# A neural circuit for flexible control of persistent behavioral states

**DOI:** 10.1101/2020.02.04.934547

**Authors:** Ni Ji, Gurrein K. Madan, Guadalupe I. Fabre, Alyssa Dayan, Casey M. Baker, Ijeoma Nwabudike, Steven W. Flavell

## Abstract

To adapt to their environments, animals must generate behaviors that are closely aligned to a rapidly changing sensory world. However, behavioral states such as foraging or courtship typically persist over long time scales to ensure proper execution. It remains unclear how neural circuits generate persistent behavioral states while maintaining the flexibility to select among alternative states when the sensory context changes. Here, we elucidate the functional architecture of a neural circuit controlling the choice between roaming and dwelling states, which underlie exploration and exploitation during foraging in *C. elegans*. By imaging ensemble-level neural activity in freely-moving animals, we identify stable, circuit-wide activity patterns corresponding to each behavioral state. Combining circuit-wide imaging with genetic analysis, we find that mutual inhibition between two antagonistic neuromodulatory systems underlies the persistence and mutual exclusivity of the opposing network states. Through machine learning analysis and circuit perturbations, we identify a sensory processing neuron that can transmit information about food odors to both the roaming and dwelling circuits and bias the animal towards different states in different sensory contexts, giving rise to context-appropriate state transitions. Our findings reveal a potentially general circuit architecture that enables flexible, sensory-driven control of persistent behavioral states.

## INTRODUCTION

The behavioral state of an animal—whether it is active, inactive, mating, or sleeping— influences its perception of and response to the environment^1–5^. In contrast to fast motor actions, behavioral states are often highly stable, lasting from minutes to hours. Despite this remarkable stability, animals can flexibly choose their behavioral state based on the sensory context and switch states when the context changes^6–8^. How the brain generates persistent behavioral states while maintaining the flexibility to select among alternative states is not well understood.

At the neural level, persistent behavioral states are often associated with stable patterns of neural activity. For example, continuous activation of pCd neurons in male Drosophila underlies persistent courtship and aggressive behaviors^9^. In addition, recent large-scale recordings of neural activity have revealed that behavioral states such as sleep and active locomotion are represented as stable, stereotyped activity patterns in neurons spanning multiple brain regions^5,6,10–13^. While the encoding of a behavioral state can be broadly distributed, the neurons that control the onset and duration of a state are often a smaller subset of those that comprise the full circuit^6,14^. To gain mechanistic insights into how persistent behavioral states are generated and controlled, it will be critical to elucidate the functional interactions among key control neurons and understand how they incorporate incoming sensory inputs that influence behavioral states.

Past studies have proposed recurrent circuitry and neuromodulation as two central mechanisms underlying persistent behavioral states. While theoretical studies have shown that recurrent excitatory or inhibitory feedback can underlie stable firing patterns^15–18^, direct experimental evidence linking recurrent circuitry with persistent activity remains scarce. Neuromodulators are known to control persistent behaviors like sleep and wake states, as well as states of stress and hunger^19–22^. However, our understanding of how ongoing neuromodulator release *in vivo* promotes persistent circuit activity remains limited. In addition, it is unclear how dynamic sensory inputs interact with recurrent circuitry and neuromodulation to elicit behavioral state transitions when the sensory environment changes.

In this study, we investigate the neural circuit mechanisms that give rise to stable, circuit-wide activity patterns during persistent foraging states in *C. elegans*. While foraging on bacterial food, *C. elegans* alternate between roaming states, characterized by sustained forward movement at high speed, and dwelling states, marked by slow movement and frequent reorientations^23,24^. Each state can last up to tens of minutes and the transitions between states are abrupt. The fraction of time an animal spends in each state is influenced by its satiety, ingestion of bacterial food, and sensory cues such as odors^23,25,26^. Consistent with the notion that these states reflect an exploration-exploitation tradeoff, animals favor dwelling in food-rich environments and after starvation, but favor roaming in poor-quality food environments and after aversive stimulation.

We and others previously found that serotonin (5-HT) and the neuropeptide pigment-dispersing factor (PDF) act as opposing neuromodulators that stabilize dwelling and roaming states, respectively^27–30^. Cell-specific genetic perturbations have uncovered the neurons that produce and detect these neuromodulators to control the stability of each behavioral state^28^. However, these identified neurons are densely interconnected with one another and with other neurons in the *C. elegans* connectome (Fig. 1B), making it impossible to infer the core functional circuitry that shapes the roaming and dwelling states from these prior studies. Crucially, it remains unclear how functional interactions between these neurons influence overall circuit activity and how 5-HT and PDF stabilize opposing states of circuit activity. In addition, while it is clear that sensory cues can influence roaming and dwelling behaviors, it remains unclear how sensory inputs converge onto this core neuromodulatory circuit to influence behavioral states.

**Figure 1.**
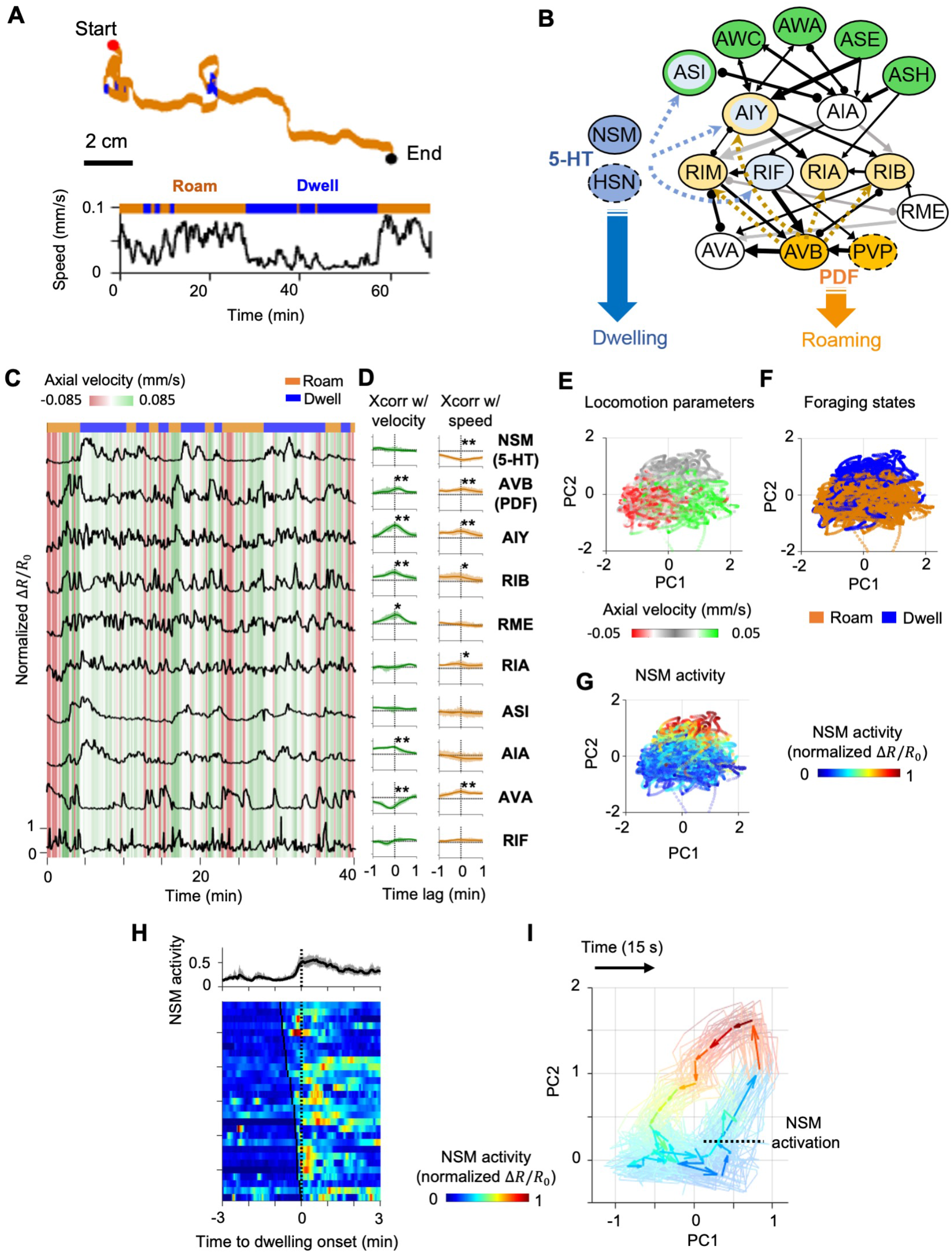
Circuit-wide calcium imaging reveals stable, low-dimensional neural representation of foraging states. (A) (Top) Movement trajectory of a *C. elegans* animal foraging on bacterial food under the tracking microscope. Red and black dots mark the beginning and end of the trajectory, respectively. Orange indicates that the animal was in the roaming state, while blue indicates the animal in dwelling state. (Bottom) The speed of the animal during the same period. (B) Putative neural circuit that mediates the sensory control of the roaming and dwelling states, based on the *C. elegans* connectome^41^ and genetic analyses from a previous study^46^. Each *C. elegans* neuron has a three-letter name. Green indicates sensory neurons. Dark blue indicates serotonergic neurons while light blue denotes neurons expressing the serotonin-gated chloride channel, MOD-1. Dark orange indicates neurons that synthesize the neuropeptide PDF, while light orange denotes neurons expressing the PDFR-1 receptor. Black arrows represent direct synaptic connections, gray arrows represent bi-synaptic connections, and lines ending in circles represent gap junctions. Thickness of these arrows indicate the number of synapses at a given connection. Dotted blue and orange arrows indicate neuromodulatory connections from Flavell et al., 2013. Neurons with dotted boundaries are located outside the head region and were not examined through calcium imaging. (C) Example dataset from multi-neuron calcium imaging in a free-moving wild-type animal. The calcium activity of each neuron is shown in black. Green-red heat map in the background indicate axial velocity of the animal, and the behavioral state of the animal is shown on top. GCaMP6m data were divided by co-expressed mScarlett fluorescence levels and normalized to the first percentile of the activity for each neuron (*R_o_*). (D) Average cross-correlation functions between individual neurons and the following locomotion variables: (left column) axial velocity, (right column) axial speed. Data are shown as mean and 95% confidence interval (95% CI). N=17 wild-type animals. *p<0.05, **p<0.01, Wilcoxon rank-sum test (E-G) Scatterplots projecting neural activity data onto principal component space, colored according to the ongoing axial velocity (E), foraging state (F), or the activity of the serotonergic neuron NSM (G). N=17 wild-type animals. (H) NSM activity aligned to the onset of dwelling states. (Top) Average NSM activity around the onset of dwelling states. (Bottom) Heat map of NSM activity around individual instances of roaming-to-dwelling transition. Dotted black line denotes the onset of dwelling states. Black ticks on the heat map mark the onset of an NSM activity bout. (I) Average circuit activity dynamics in principal component space aligned to the onset of NSM activation. Each colored arrow represents average activity dynamics over a 15 second interval. Color indicates ongoing NSM activity. Faint lines show bootstrap samples of the average dynamics.

To address these questions, we performed simultaneous calcium imaging of defined neurons throughout the roaming-dwelling circuit in freely-moving animals. We identified stereotyped, circuit-wide activity patterns corresponding to each foraging state. By combining circuit imaging with genetic perturbations, we identified a mutual inhibitory loop between the serotonergic NSM neuron and the 5-HT and PDF target neurons. We found that this mutual inhibition is critical for the persistence and mutual exclusivity of the circuit-wide activity states that correspond to roaming and dwelling. Through machine learning analyses and circuit perturbations, we found that the AIA sensory processing neuron sends parallel outputs to both neuromodulatory systems and can bias the network towards either roaming or dwelling, depending on the sensory context. Together, these results identify a functional circuit architecture that allows for flexible, sensory-driven control of persistent behavioral states.

## RESULTS

### Roaming and dwelling states are associated with stable, circuit-wide activity changes

To understand how roaming and dwelling states arise from circuit-level interactions between neurons, we sought to monitor the activity of neurons throughout the core roaming-dwelling circuit in wild-type animals and in mutant backgrounds that perturb signaling among the neurons. We built a calcium imaging platform with a closed-loop tracking system that allows for simultaneous imaging of many neurons as animals freely move (Fig. 1A and Figure 1-Figure Supplement 1A-B)^31–33^. We generated a transgenic line where well-defined promoter fragments were used to express GCaMP6m in a select set of 10 neurons (Fig. 1B and Figure 1-Figure Supplement 1C). These neurons were selected based on their classification into at least one of the three groups: 1) neurons expressing 5-HT, PDF, or their target receptors MOD-1 or PDFR-1^28^, 2) neurons that share dense synaptic connections with those in 1), and 3) premotor or motor neurons whose activities report the locomotory intent of animals^34,35^. We performed circuit-level imaging of these animals as they foraged on uniformly-seeded bacterial lawns. Imaging this defined subset of neurons in many animals allowed us to leverage prior knowledge and easily determine the identity of each neuron in each recording, thereby circumventing the challenge of determining neuronal identity in a densely-labeled brain.

Because roaming and dwelling are characterized by marked changes in locomotion, we first asked how locomotion parameters, such as movement direction and speed, are encoded by circuit activity. Six of the recorded neurons displayed calcium signals that co-varied with the animals’ movement direction (Fig. 1C and 1D, right; see correlation with axial velocity). These include the PDF-1-expressing neuron AVB and PDFR-1-expressing neurons AIY and RIB, all known to promote forward runs^34–37^, as well as the premotor neuron AVA, known to promote reversals^35,38^. In addition, several neurons exhibited calcium signals that were correlated with locomotion speed. The serotonergic neuron NSM was most active at low speeds, while roaming-promoting neurons such as AVB and AIY were most active at high speeds (Fig. 1C and 1D). These data suggest that locomotion parameters of the animal are encoded by neurons distributed throughout the roaming-dwelling circuit.

The encoding of direction and speed across many neurons suggests a circuit-wide representation of the animal’s behavioral state. To test whether the dominant modes of activity in the circuit were associated with the animal’s behavioral state, we performed Principle Component Analysis (PCA) using the activity profiles of all the recorded neurons and examined whether the top PCs were associated with roaming and dwelling (Fig. 1E-F). Indeed, the top two principle components (PC1 and PC2), which together explain 44% of the total variance, exhibited clear behavioral correlates. While neural activity during forward and reverse locomotion segregated along PC1 (Fig. 1E), roaming and dwelling states corresponded to low and high values on PC2 (Fig. 1F). Our observation that PC1 encodes forward-reverse movement matches previous population recordings from immobilized and non-feeding animals^11^. However, our finding that PC2 represents roaming-dwelling states is notably different from these prior studies. These results suggest that the *C. elegans* neural activity space is constrained in some respects, but also varies considerably across different environmental conditions. Overall, these results indicate that the main sources of activity variance in the circuit are associated with rapid locomotion dynamics (PC1) and stable foraging states (PC2). This robust mapping between circuit activity and behavior suggests that persistent circuit dynamics may underlie stable roaming and dwelling states.

### Persistent NSM activation and associated circuit-wide activity changes correspond to the dwelling state

To understand how the activities of individual neurons were related to these dominant modes of dynamics in the circuit, we next examined the loadings of neurons on PC1 and PC2 (Figure 1-Figure Supplement 3B). Several of the locomotion-encoding neurons were strongly represented on both PC1 and PC2, suggesting that each of these neurons’ activities reflect forward-reverse locomotion, as well as the animal’s foraging state. In contrast, the serotonergic neuron NSM was strongly represented in PC2, but was largely absent in PC1. Consistent with this, we found that NSM activity co-varied with PC2 values, but not with PC1 (Fig. 1G). This suggests that NSM activity changes as animals switch between foraging states, but not as they rapidly switch between forward and reverse locomotion. NSM activation was remarkably persistent, closely paralleling the duration of dwelling states: the onset of NSM activity was reliably followed by a rapid drop in speed and the onset of dwelling, while its offset was frequently accompanied by an increase in speed and the transition back to roaming (Fig. 1H and Figure 1-Figure Supplement 4A-C). Together with previous work that showed that optogenetic NSM activation can drive animals into dwelling states via its release of serotonin^28,39^, these observations indicate that NSM activation is closely tied to the dwelling state and may play an important role in organizing the circuit-wide activity state that corresponds to dwelling.

To further explore this possibility, we examined how circuit activity evolved in PC space during periods of NSM activation. We found that NSM activation often began when circuit activity was in the region of the PC space with high PC1 activity and low PC2 activity, typical of forward locomotion (Fig. 1I). As NSM became active, circuit activity rose rapidly along PC2 (each arrow in Fig. 1I represents 15 sec). After reaching a peak on PC2, circuit activity slowly traveled towards low values of both PC1 and PC2, hitting a corner of the PC space corresponding to the animal in reverse locomotion. Optogenetic activation of NSM in roaming animals evoked a similar trajectory in circuit activity, starting from high PC1 and low PC2 regions of the PC space and progressing in a counter-clockwise fashion (Figure 1-Figure Supplement 4D). These results suggest that the persistent activation of NSM during dwelling is associated with stereotyped changes in overall circuit dynamics.

### Persistent activity in serotonergic NSM neurons requires feedback from its target neurons that express the MOD-1 serotonin receptor

The above analyses of wild-type circuit dynamics suggest that stereotyped circuit-wide activity patterns are associated with roaming and dwelling states. We next examined how these neural dynamics are influenced by neuromodulatory connections embedded in the circuit. Although the 5-HT and PDF systems are known to act in opposition to regulate roaming and dwelling behaviors^28^, it is not known how these neuromodulators impact circuit dynamics. To address this, we imaged neural activity in mutants deficient in 5-HT signaling, PDF signaling, or both (Fig. 2 and 3). Mutants that disrupt 5-HT signaling, such as those lacking a key enzyme for serotonin biosynthesis (*tph-1*) or a 5-HT-gated chloride channel (*mod-1*), exhibited a decrease in time spent in the dwelling state (Fig. 2A-C). Interestingly, we found that the durations of the long-lasting NSM activity bouts were also dramatically shortened in these mutants, resulting in a significant decrease in overall NSM activity (Fig. 2D-E). This result indicates that 5-HT signaling is required to sustain the activity of the serotonergic neuron NSM. Because MOD-1 is an inhibitory 5-HT-gated chloride channel, these data indicate that the *mod-1*-expressing neurons must play an inhibitory role in regulating NSM activity. Previous work has shown that *mod-1* functions in the neurons AIY, RIF, and ASI to promote dwelling^28^ (Fig. 1B). Since none of these neurons directly synapse onto NSM, they must functionally inhibit NSM through a polysynaptic route or via the release of a neuromodulator. Together, these results indicate that the serotonergic NSM neuron reinforces its own activity via mutual inhibition with neurons expressing the inhibitory 5-HT receptor MOD-1 (Fig. 2F).

**Figure 2.**
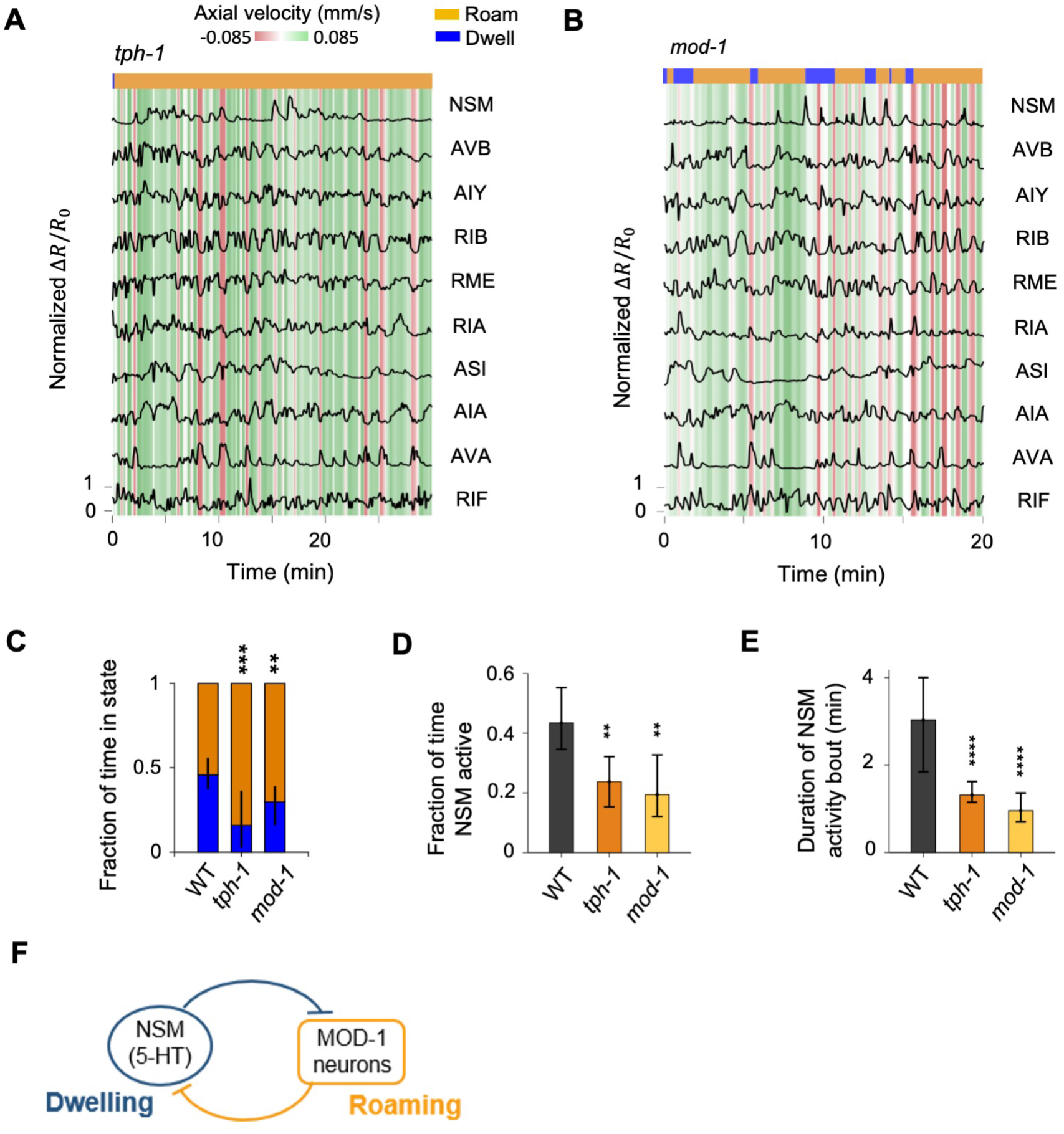
Serotonin signaling promotes persistent activation of serotonergic NSM neurons via a mutual inhibitory circuit. (A-B) Example circuit-wide calcium imaging datasets from *tph-1* (A) and *mod-1* (B) mutant animals, shown as in Fig. 1C. (C) Fraction of time animals spent roaming versus dwelling for wild-type, *tph-1*, and *mod-1* animals. Error bars depict mean and 95% CI. (D) Fraction of time NSM is active for animals of the indicated genotypes. Data are shown as mean and 95% CI. (E) Duration of NSM activation bouts for the indicated genotypes, shown as mean and 95% CI. For (C-E), N = 17, 10, and 8 animals for WT, *tph-1*, and *mod-1*, respectively. (F) Circuit schematic based on results from the *tph-1* and *mod-1* mutants, showing cross inhibition between the 5-HT neuron NSM and the MOD-1 expressing neurons. **p<0.01, ***p<0.001, ****p<0.0001, Wilcoxon rank-sum test with Benjamini-Hochberg (BH) correction.

**Figure 3.**
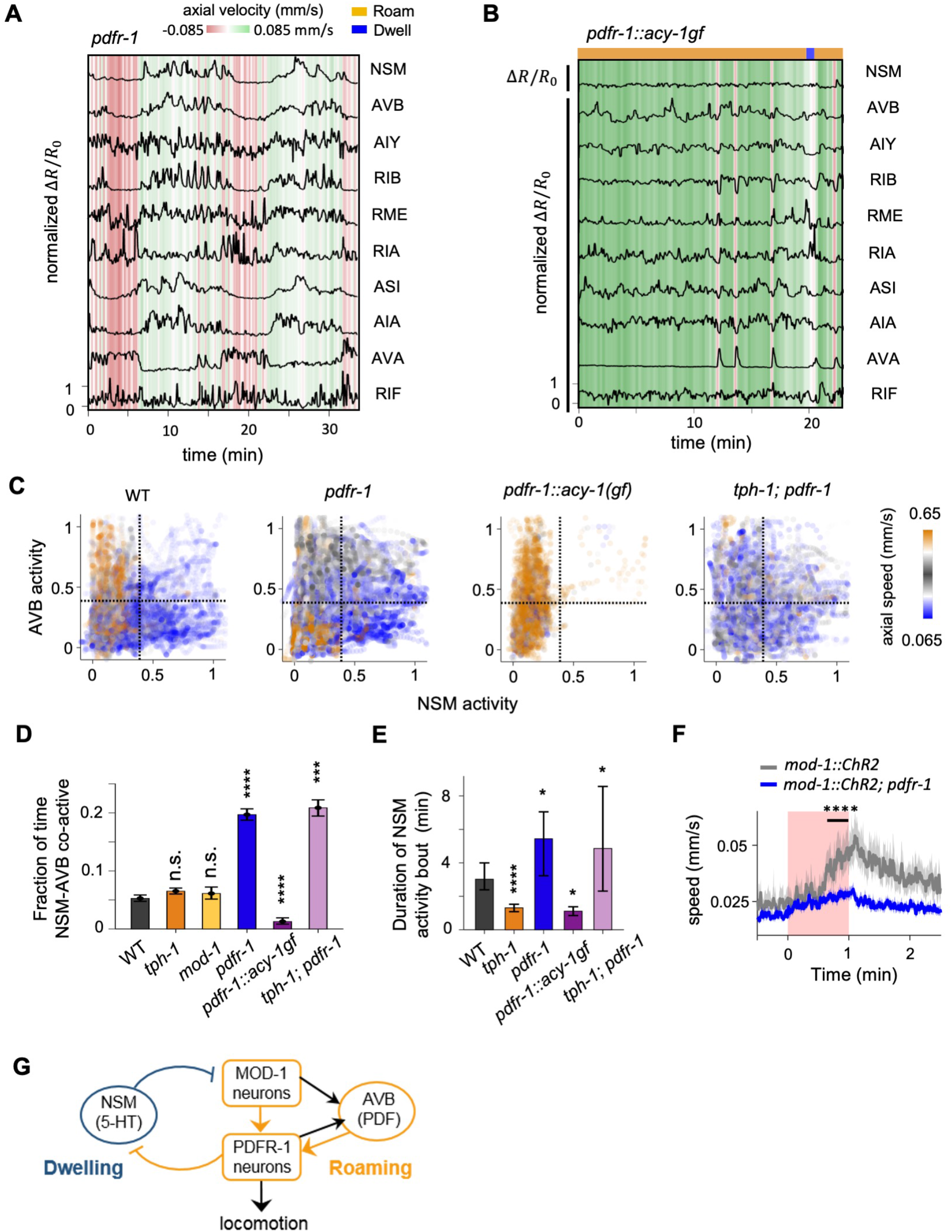
PDF signaling is required for mutual exclusivity between circuit states and acts downstream of the 5-HT target neurons in the mutual inhibitory circuit. (A) Example circuit-wide calcium imaging dataset from *pdfr-1* mutants lacking PDF neuropeptide signaling, shown as in Fig. 1C. No roaming/dwelling ethogram is shown for *pdfr-1* animals due to changes in their speed distribution that implicate altered or new behavioral states (see Figure 3-Figure Supplement 2B-C). (B) Example circuit-wide calcium imaging dataset from transgenic animals expressing the hyperactive PDFR-1 effector ACY-1(P260S) specifically in PDFR-1 expressing neurons. For NSM, the un-normalized Δ*R*/*R_o_* is shown since the Δ*R*/*R_o_* values never exceeded 10% of the average peak NSM activity wild-type animals. (C) Scatterplots of NSM and AVB activity in *pdfr-1* mutants, transgenic *pdfr-1::acy-1(P260S)gf* animals, and *tph-1; pdfr-1* double mutants. Data points are colored by the instantaneous speed of the animal. Color scale was chosen so that blue colors correspond to speeds typical of the dwelling state, orange correspond to speeds typical of the roaming states, while gray colors indicate intermediate speeds. (D) Fraction of time in which both NSM and AVB are active. ***p<0.001; ****p<0.0001, bootstrap estimates of the mean with BH correction. (E) Duration of NSM activity bouts in the indicated genotypes shown as mean and 95% CI. *p<0.05; ****p<0.0001, Wilcoxon rank-sum test with BH correction. For (C-E), N = 17, 10, 8, 11, 9 and 8 animals for WT, *tph-1*, and *mod-1*, *pdfr-1*, *pdfr-1::acy-1gf*, and *tph-1;pdfr-1* animals. The data for *tph-1* animals are the same as in Fig. 2E. (F) Speed of wild-type and *pdfr-1* mutant animals in response to optogenetic activation of the MOD-1 expressing neurons (red shading). Average speeds during the window spanned by the black line were compared between animals of the two genotypes. ****p<0.0001, Wilcoxon rank-sum test. (G) Circuit schematic summarizing results shown in (C-G): the PDFR-1 expressing neurons act downstream of the MOD-1 expressing neurons to inhibit the 5-HT neuron NSM. Black arrows indicate anatomical connections based on the *C. elegans* connectome^51^.

### PDF receptor-expressing neurons inhibit NSM to promote mutually-exclusive circuit states

We next examined the impact of PDF signaling on circuit dynamics by imaging animals carrying a null mutation in the PDF receptor gene *pdfr-1* (Fig. 3A). In wild-type animals, the serotonergic neuron NSM and the PDF-1-producing neuron AVB exhibited a mutually exclusive activity pattern corresponding to the roaming and dwelling states (Fig. 3C). This mutual exclusivity between NSM and AVB was disrupted in *pdfr-1* mutants (Fig. 3C-D). In these mutant animals, the two neurons were frequently co-active, giving rise to a positive correlation between the activities of the two neurons (Fig. 3D and Figure 3-Figure Supplement 1C). Positive correlations also appeared between NSM and other roaming-active neurons, including the *pdfr-1*-expressing neurons AIY and RIB (Figure 3-Figure Supplement 1C). Interestingly, we observed that *pdfr-1* animals frequently moved at speeds mid-way between those typically seen for roaming and dwelling states in wild-type animals (Figure 3-Figure Supplement 2B-C). Thus, ectopic co-activation of the dwelling-active NSM neuron and the roaming-active neurons likely results in behavioral outputs that blur the boundary between roaming and dwelling. Together, these findings indicate that PDF signaling is required for the neural circuit to maintain mutual exclusivity between the opposing circuit states that underlie roaming and dwelling.

In contrast to the *tph-1* animals, NSM activity bouts in *pdfr-1* mutants were more persistent than they were in wild-type animals (Fig. 3E). This result suggests that PDF signaling plays an important role in suppressing NSM activity. Consistent with this interpretation, constitutive activation of PDFR-1 signaling, via expression of the hyperactive PDFR-1 effector ACY-1(P260S) in the *pdfr-1*-expressing neurons, strongly inhibited NSM activity (Fig. 3B-E). Together, these findings indicate that PDF signaling in the PDFR-1-expressing neurons is necessary and sufficient to keep NSM inactive during roaming, a key requirement for maintaining mutual exclusivity between the roaming and dwelling states.

### A mutual inhibitory loop links serotonergic NSM neurons with the MOD-1- and PDFR-1- expressing neurons

To probe whether the MOD-1- and PDFR-1-expressing neurons act in the same pathway to suppress NSM activity, we performed epistasis analysis by examining *tph-1;pdfr-1* double mutants. Similar to *pdfr-1* mutants, these animals exhibited ectopic co-activity of NSM and AVB and prolonged bouts of NSM activation, suggesting that *pdfr-1* functions downstream of *tph-1* to control NSM activity (Fig. 3C-E and Figure 3-Figure Supplement 2A). Consistent with this observation, we also found that optogenetic activation of the *mod-1*-expressing neurons, which triggered roaming in wild-type animals, failed to do so in *pdfr-1* mutants (Fig. 3F). *pdfr-1* is not expressed in NSM and functions in multiple other neurons, such as RIM, AIY, RIA, and RIB, to promote roaming^39,40^. Thus, these data suggest that the PDFR-1-expressing neurons act downstream of the MOD-1-expressing neurons to inhibit NSM activity (Fig. 3G).

Altogether, these results indicate that mutual inhibition between NSM and the neurons that express the MOD-1 and PDFR-1 receptors is necessary for the stability and mutual exclusivity of the circuit states corresponding to roaming and dwelling. Based on the *C. elegans* connectome^41^ and previous studies^34,42–45^, many of the MOD-1- and PDFR-1-expressing neurons receive synaptic inputs from sensory neurons and are functionally involved in sensorimotor behaviors (Fig. 1B). The prominent positions of these neurons in sensory processing circuits raise the possibility that they may serve important roles in conveying information about sensory cues to the roaming-dwelling circuit.

### A CNN classifier reveals stereotyped circuit dynamics that precede roaming-to-dwelling transitions

The functional circuit architecture revealed through our calcium imaging analyses raised the possibility that incoming sensory inputs that act on the MOD-1- and PDFR-1-expressing neurons might influence the transitions between roaming and dwelling. One prediction of this hypothesis is that these neurons that receive sensory inputs may display reliable activity patterns prior to state transitions.

To test the above hypothesis, we sought to predict state transitions from prior circuit activity dynamics. Our calcium imaging results showed that the onset of NSM activity reliably coincided with the onset of dwelling states (Fig. 1H). This observation, combined with prior studies showing that serotonin release from NSM triggers roaming-to-dwelling transitions^46^, suggests that NSM activation is a key event that drives the transition to dwelling. We thus focused on uncovering potential circuit elements that function upstream of NSM to drive the roaming-to-dwelling transition. We adopted a supervised machine learning approach by training a Convolutional Neural Network (CNN) classifier to predict NSM activation using the preceding multi-dimensional activity profile from all other neurons imaged (Fig. 4A-B; see Methods). We chose the CNN classifier because of its flexible architecture, which can model complex nonlinear relationships between the input and output variables and detect multiple relevant activity patterns via the same network^47–49^. Successfully trained networks achieved over 70% test accuracy, equaling or exceeding other supervised learning methods (Figure 4-Figure Supplement 1A). This result indicates that stereotyped circuit activity patterns frequently precede NSM activation.

**Figure 4.**
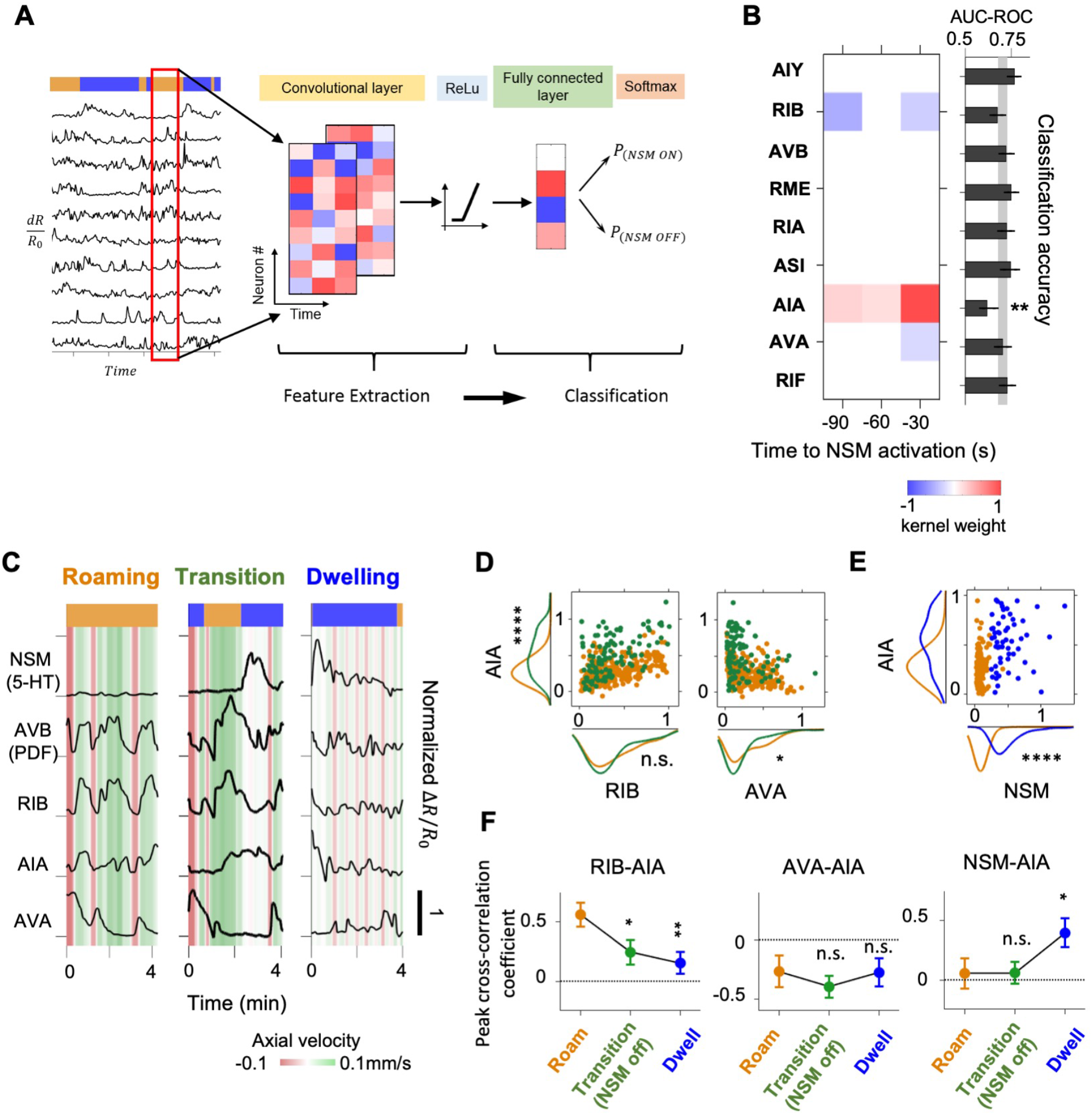
A CNN classifier identifies circuit activity patterns predictive of roaming-to-dwelling state transitions. (A) Schematic illustrating the architecture of the Convolutional Neural Network (CNN) trained to predict NSM activation events. (B) Left: a common convolutional kernel found across successfully trained CNNs. Only weights that are significantly different from zero are colored. Right: Feature selection results. Each black bar depicts the average area under the curve for the Receiver Operating Characteristic curve (AUC-ROC) from networks trained using data with one neuron held out at a time. The identity of the held-out neuron is indicated to the far left. The gray stripe in the background denotes the 95% CI of the AUC-ROC from networks trained using data from all 9 neurons. Error bars are 95% CI of the mean. **p<0.01, bootstrap estimate of the mean with BH correction. (C) Example activity traces from NSM, AVB, and the three neurons with significant weights in the convolutional kernel. Activity traces were taken during roaming (left), dwelling (right) and roaming-to-dwelling transition. (D) Scatterplots of simultaneously measured neural activity of AIA and RIB (left) or AIA and AVA (right). Orange data points are taken during roaming states at least 1 minute before the onset of the next dwelling states and before NSM becomes active. Green data points are taken within 1 minute before the onset of dwelling states. Along the x- and the y-axes are marginal probability distributions of the data points shown in the scatterplots. (E) Scatterplots of simultaneously measured neural activities of AIA and NSM. Orange data points are taken within 1 minute before the onset of the next dwelling states and before NSM becomes active. Blue data points are taken within 30 seconds after the onset of dwelling states. Along the x- and the y-axes are marginal probability distributions of the data points shown in the scatterplots. (F) Peak cross-correlation coefficients among the key neurons during roaming (data taken from 100-70 seconds before the onset of the next dwelling state), roaming-to-dwelling transition (data taken from 30-0 second before the NSM activation event prior to the onset of the next dwelling state), or dwelling (data taken from 10-40 seconds after dwelling onset). See Figure 4-Figure Supplement 2 for full cross-correlation profiles. Error bars are standard error of the mean. For (B-F), N = 17 WT animals. For (D-F), *p<0.05, **p<0.01, ****p<0.0001, Wilcoxon rank-sum test with BH correction.

We examined the parameters of the trained networks to further define the activity patterns that were being used to make successful predictions about upcoming NSM activation. Successfully trained networks consistently employed a convolutional filter where the largest positive weights were associated with the sensory processing neuron AIA and the largest negative weights were linked to two locomotion-promoting neurons RIB and AVA (Fig. 4B). These weights suggest that NSM activation is most likely to occur following heightened activity in AIA and attenuated activity in RIB and AVA. Withholding AIA, RIB, and AVA from the training data abolished the predictive power of the trained network, while withholding AIA activity alone also led to a significant reduction in test accuracy (Fig. 4B and Figure 4-Figure Supplement 1B). Moreover, networks trained on the activities of only AIA, RIB, and AVA performed nearly as well as those trained on all the neurons (Figure 4-Figure Supplement 1B). These observations suggest that the combined activities of AIA, RIB, and AVA can frequently predict the onset of NSM activity.

We next examined how the activities of AIA, RIB and AVA changed during transitions from roaming to dwelling. During roaming, AIA activity was positively correlated with that of forward run-promoting neurons, such as RIB, and negatively correlated with the reversal-promoting neuron AVA (Fig. 4C-E; Figure 4-Figure Supplement 2). Within 30 seconds prior to NSM activation, AIA often exhibited a further increase in activity, while RIB and AVA activity stayed at similar levels or decreased. As NSM activity rose and the animal entered the dwelling state, RIB and AVA activity declined sharply while AIA activity became correlated with NSM. AIA then declined to baseline over the following minutes. Thus, AIA activity co-varies with the forward-active neurons during roaming and with NSM at the onset of dwelling (Fig. 4F). This native activity pattern is consistent with the convolutional kernel from the CNN classifier, where heightened activity of AIA relative to the locomotion-promoting neurons RIB and AVA predicts NSM activation. Together, these results reveal a stereotyped, multi-neuron activity pattern that predicts NSM activation.

### AIA activation can elicit both roaming and dwelling states

Because AIA activity co-varied with both roaming- and dwelling-active neurons and was required for the prediction of NSM activation, we hypothesized that AIA might play an active role in controlling the transitions between roaming and dwelling. To test this, we optogenetically activated AIA in foraging animals exposed to uniform lawns of bacterial food (Fig. 5A-B). Behavioral responses to AIA activation depended on the state of the animal at the time that AIA was activated. Animals that were roaming at that time had opposite behavioral responses to those that were dwelling. Roaming animals on average exhibited a rapid and transient decrease in speed upon AIA activation, while dwelling animals showed a gradual increase in speed upon AIA activation (Fig. 5A-B). Among the roaming animals, ∼40% entered dwelling within seconds of stimulation onset. By 40 seconds into the stimulation, however, most of the animals that were roaming pre-stimulation either remained in or re-entered the roaming state. Among animals that were dwelling pre-stimulation, no difference was observed between the optogenetically-stimulated and control animals until 40 seconds after stimulation onset, at which time ∼30% of animals had transitioned into the roaming state. These results indicate that optogenetic activation of AIA can affect state transitions on two different time scales: triggering the roaming-to-dwelling transition within a few seconds and promoting entry into the roaming state upon tens of seconds of continued activation.

**Figure 5.**
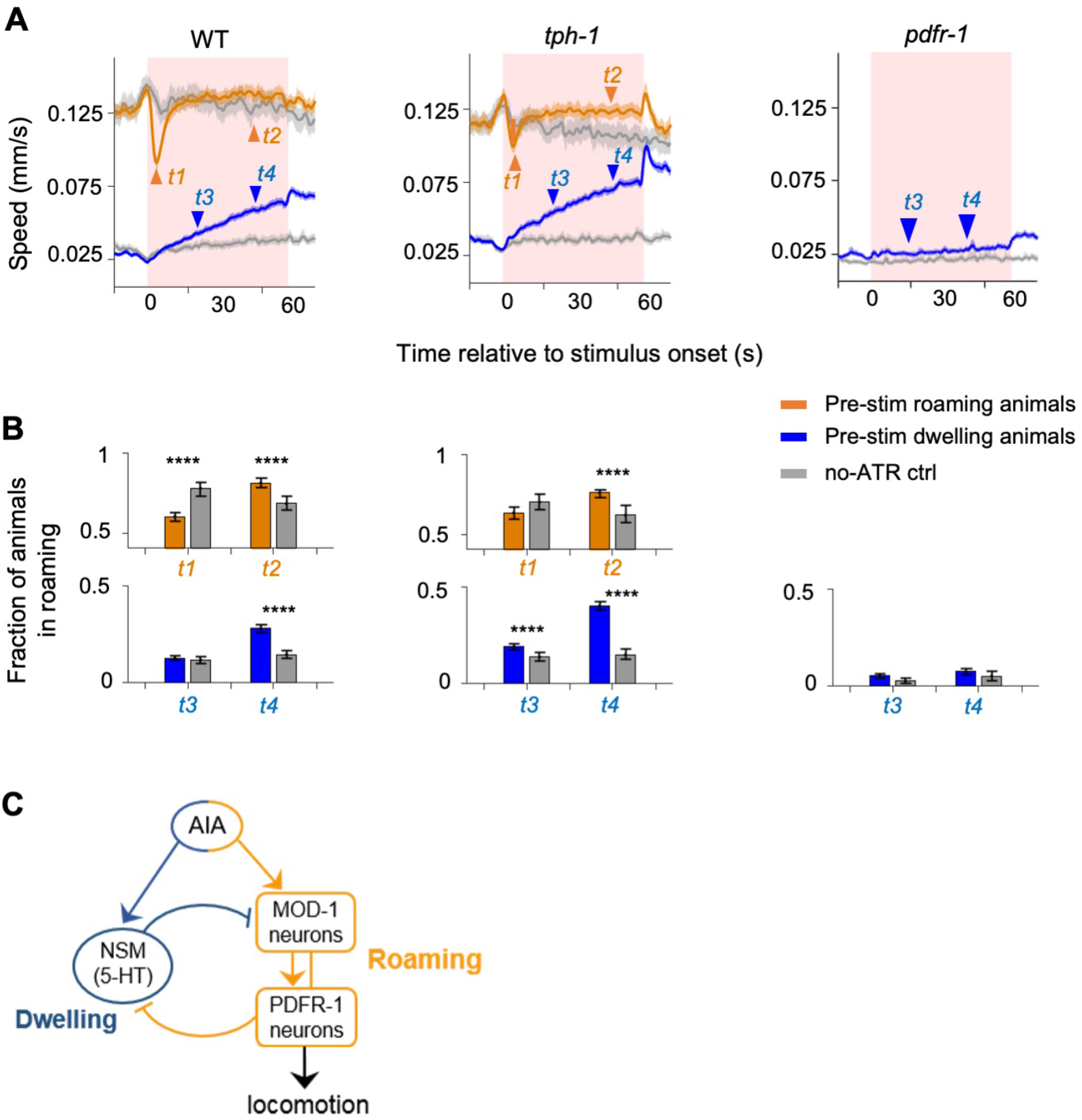
The AIA sensory processing neuron can drive behavioral state switching. (A) Average locomotion speed before, during, and after AIA::Chrimson activation for wild-type (left), *tph-1* (middle), and *pdfr-1* (right) animals. Animals were grouped by whether they were roaming (orange) or dwelling (blue) prior to AIA stimulation. Pink patches in the background denote the one-minute stimulation window. Gray lines indicate no-all-trans-retinal (no-ATR) controls. N=1032 wild-type animals were compared to N=370 no-ATR controls. N= 927 *tph-1* mutants were compared to N = 284 no-ATR control. N= 383 *pdfr-1* mutants were compared to N = 237 no-ATR controls. Note that roaming in *pdfr-1* animals was too rare and brief to be included for analysis on AIA-induced slowing. Orange and blue arrowheads denote time points used for analyses in (B). Error bars are 95% CI of the mean. (B) Fraction of animals in the roaming state at different phases of AIA::Chrimson stimulation. Top: Among animals that were roaming pre-stimulation, the fraction of them that were roaming after 4 seconds or 40 seconds from the onset of AIA stimulation. Bottom: Among animals that were dwelling pre-stimulation, the fraction of them that were roaming after 20 seconds or 40 seconds from the onset of AIA stimulation. Same analyses were performed for wild-type (left), *tph-1* (middle), and *pdfr-1* (right) animals. See panel (A) for full traces. Error bars are 95% CI of the mean. **** p<0.0001, Wilcoxon rank-sum test. (C) Functional architecture of the circuit controlling the roaming and dwelling states, based on results from Figures 2-5.

To test whether AIA acts through 5-HT and PDF to drive these observed changes in behavior, we optogenetically activated AIA in mutants defective in 5-HT or PDF signaling (*tph-1* and *pdfr-1* animals, Fig. 5A-B). In *tph-1* mutants, animals that were roaming pre-stimulation no longer displayed rapid entry into dwelling and showed a higher probability of returning to roaming later into the stimulation. *tph-1* mutants that were dwelling pre-stimulation displayed a higher probability of entering the roaming state during stimulation. Conversely, AIA activation in *pdfr-1* mutants that were dwelling pre-stimulation failed to elicit transitions into roaming. Roaming states in these mutants were too infrequent and brief to warrant meaningful analysis. Together, these results indicate that AIA promotes dwelling via 5-HT signaling and promotes roaming via PDF signaling. Because the 5-HT-dependent slowing response can be independently perturbed from the PDFR-1-dependent speeding response, these results raise the possibility that AIA provides parallel outputs to both neuromodulatory systems (Fig. 5C).

### AIA can promote either roaming or dwelling, depending on the sensory context

Based on the *C. elegans* connectome^50,51^, AIA receives the majority of its synaptic inputs (∼80%) from chemosensory neurons (Figure 4-Figure Supplement 1C), many of which detect temporal changes in the concentrations of olfactory and gustatory cues^52–54^. Previous work has shown that AIA is activated by an increase in the concentration of attractive odorants present in bacterial food^53,55^. In the absence of food, AIA promotes forward runs when animals detect increases in attractive odors^53^. AIA sends synaptic output to multiple neurons in the sensorimotor pathway, including several *mod-1-* and *pdfr-1-*expressing neurons, though its role in roaming and dwelling behaviors has not been explored.

Based on AIA’s established role in sensory processing and our observations that AIA could drive both roaming- and dwelling-like behaviors, we hypothesized that AIA may promote either roaming or dwelling, depending on the sensory cues in the environment. To test this hypothesis, we examined the foraging behaviors of wild-type animals in different sensory contexts, and compared them to animals in which AIA had been silenced (*AIA::unc-103gf*). Given that AIA responds to food odors, we developed a patch foraging assay in which animals placed on a sparse food patch can navigate a food odor gradient to approach an adjacent dense food patch (Fig. 6A). This assay is notably different from standard chemotaxis assays, where animals are not in contact with any food source and therefore do not display roaming or dwelling behaviors. To examine AIA’s impact on roaming and dwelling in the absence of an olfactory gradient, we performed a second assay where wild-type or AIA-silenced animals were presented with uniform-density bacterial food, similar to the above experiments (Fig. 6G).

**Figure 6.**
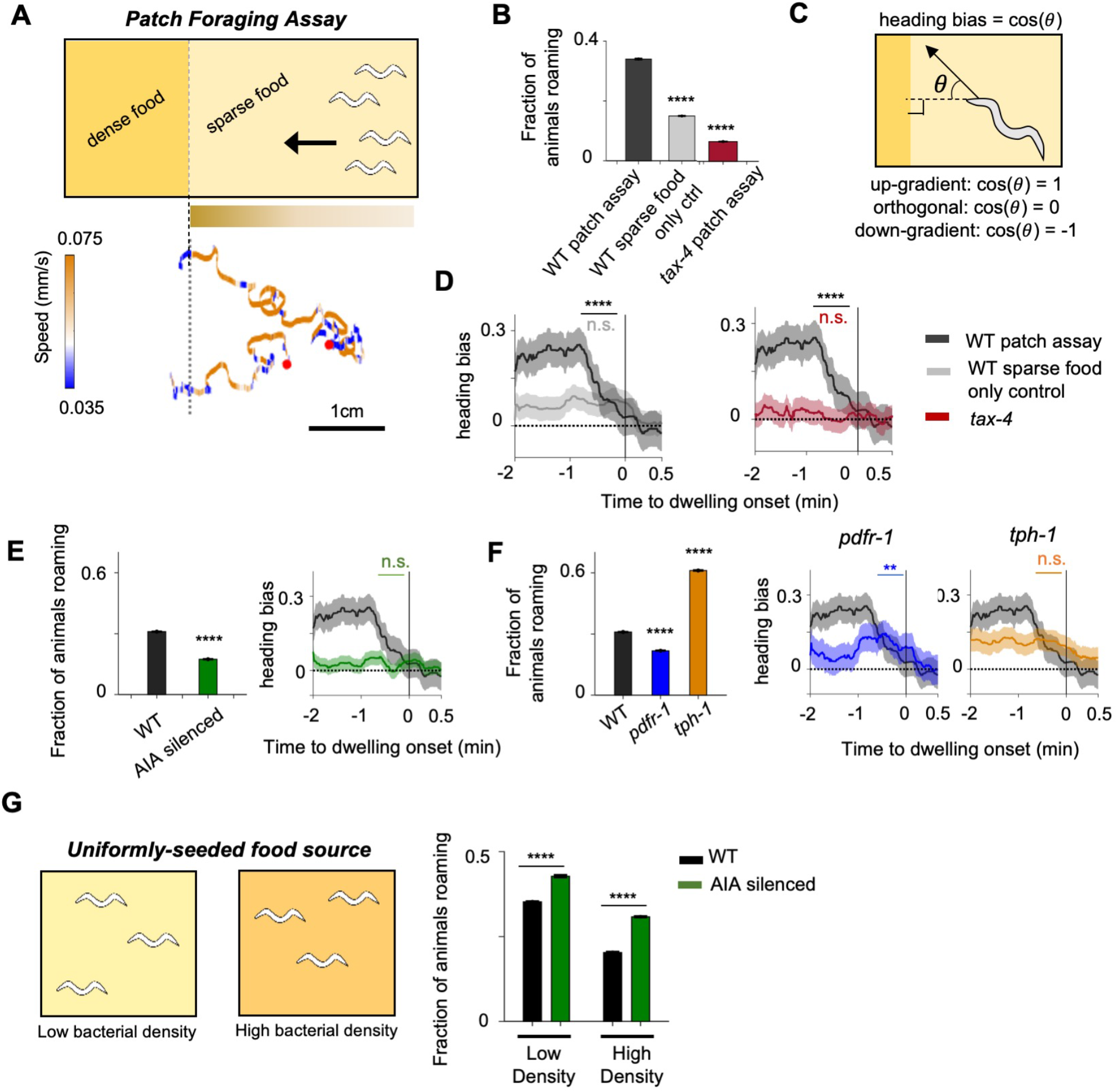
The AIA sensory processing neuron can promote either roaming or dwelling, depending on the sensory context. (A) Top: Cartoon depicting the patch foraging behavioral assay. Horizontal bar with gradient signifies the food odor gradient emanating from the dense food patch. Bottom: example trajectories of two animals from a patch foraging assay. Color scale indicates speed, with orange corresponding to roaming-like speeds and blue dwelling-like speeds. Red dots denote the starting points of the animals. (B) Average fraction of animals roaming on the sparse food patch in the patch foraging assay. Comparisons are made between wild-type animals in the patch foraging assay (n=288), wild-type animals assayed on uniform sparse food with no dense patch around (n=194), and *tax-4* animals in the patch foraging assay (n=81). (C) Schematic depicting how heading bias is calculated. (D) Event-triggered averages showing average heading bias of animals for two minutes prior to transitions into dwelling states. Experimental conditions are depicted with same color scheme as in (B). Data are shown as means ± SEM. The average heading bias within two time windows, one from 60-50 seconds prior to dwelling onset, the other from 20-10 seconds prior to dwelling onset, were compared. (E) Left: Average fraction of animals roaming on sparse food in the patch foraging assay in wild-type (black) versus AIA silenced (*AIA::unc-103gf*) animals (green). Right: Heading bias of AIA-silenced animals (green) two minutes prior to the transition into the dwelling state. n=197. Wild type data (black with gray error bar) are shown for comparison. (F) Left: Average fractions of animals roaming on the sparse food patch for *pdfr-1* (blue) and *tph-1* (orange) mutant animals during the patch foraging assay. Right: Heading bias two minutes prior to the transition into the dwelling state for *pdfr-1* (blue) and *tph-1* (orange) mutant animals. n=99 for *pdfr-1* and n=212 for *tph-1* animals. Wild type data (black with gray error bar) are shown for comparison. (G) Left: schematic of behavioral assays in uniformly-seeded food environments. Right: Average fractions of animals roaming for wild-type (black bars) and AIA-silenced (green bars) animals exposed to two different densities of uniformly-distributed sparse food. For all calculations on fraction of animals roaming, error bars are 95% CI of the mean. For all calculations of heading bias, error bars are SEM. For all comparisons, **p<0.01, ****p<0.0001, Wilcoxon rank sum test with BH correction.

In the patch foraging assay, wild-type animals exhibited directed motion towards the dense food patch and alternated between roaming and dwelling as they approached (Fig. 6A, bottom). Compared to control plates without the dense food patch, animals in the patch foraging assay spent more time in the roaming state (Fig. 6B), and biased their movement towards the dense food patch as they roamed (Figure 6-Figure Supplement 1A). Animals preferentially switched from roaming to dwelling when their direction of motion (measured as heading bias; Fig. 6C) began to deviate away from the dense food patch (Fig. 6D). Because the animal’s heading direction directly impacts the change in odor concentration that it experiences, these results indicate that dynamic changes in the concentration of food odors influences the transition rates between roaming and dwelling. Supporting this interpretation, we found that chemosensation-defective *tax-4* mutants^56^ subjected to the patch foraging assay failed to exhibit elevated roaming and failed to couple the roaming-to-dwelling transition with their direction of motion (Fig. 6B, 6D).

We next asked whether AIA was necessary for the sensory-induced modulation of roaming and dwelling states in the patch foraging assay. We found that AIA-silenced animals (*AIA::unc-103gf*) exhibited an overall decrease in roaming compared to wild-type animals and did not selectively enter dwelling states when their movement direction deviated away from the dense food patch (Fig. 6E and Figure 6-Figure Supplement 1B). These results indicate that AIA is necessary for animals to display elevated roaming in the presence of a food odor gradient and for animals to couple their movement direction with roaming-to-dwelling transitions.

We also examined the roles of 5-HT and PDF in the patch foraging assay. We found that *pdfr-1* mutants failed to elevate roaming in the odor gradient but were still able to couple the roaming-to-dwelling transition to their direction of motion (Fig. 6F and Figure 6-Figure Supplement 1B). In contrast, *tph-1* mutants displayed enhanced roaming but did not couple the roaming-to-dwelling transition to their direction of motion (Fig. 6F and Figure 6-Figure Supplement 1B). These results are consistent with a circuit architecture where each neuromodulatory system independently receives sensory inputs to drive the behavioral state that it controls.

Lastly, to examine the necessity of AIA for roaming and dwelling in the absence of a strong sensory gradient, we compared the behavior of wild-type and AIA-silenced animals in environments with uniformly-seeded bacterial food. We tested two different bacterial densities (Fig. 6G). In both cases, AIA-silenced animals displayed a significant decrease in the fraction of time spent dwelling (Fig. 6G). These results suggest that AIA functions to promote the dwelling state in a constant sensory environment, reminiscent of its function in promoting dwelling when animals travel orthogonal to the odor gradient in the patch foraging assay. Taken together with the above data, these results indicate that AIA can promote either roaming or dwelling, depending on the sensory environment. Overall, these data suggest that AIA plays a critical role in the sensory-dependent modulation of roaming and dwelling states.

### A minimal neural circuit model suggests that AIA’s functional impact on roaming and dwelling may depend on the exact combination of sensory inputs

To gain a better understanding of how the functional architecture of the roaming-dwelling circuit allows AIA to promote different states in different sensory contexts, we built a computational model where a three-node neural circuit dictates the behavioral state of a simulated agent navigating virtual environments (Fig. 7A-B). The topology of the model circuit recapitulates the functional architecture of the biological circuit (Fig. 7A): 1) dynamic olfactory cues are channeled through the “AIA” neuron, which sends parallel outputs to both the “5-HT” neuron and the “PDFR” neuron; 2) the “5-HT” and “PDFR” neurons mutually inhibit one another; and 3) food ingestion-related cues directly activate the “5-HT” neuron and are required for its activity^39^. Depending on the relative activities of the “5-HT” and “PDFR” neurons in the circuit model, the agent will either be in a paused “dwelling state” or a fast moving “roaming state,” in which chemotaxis takes place through klinotaxis (see Methods).

**Figure 7.**
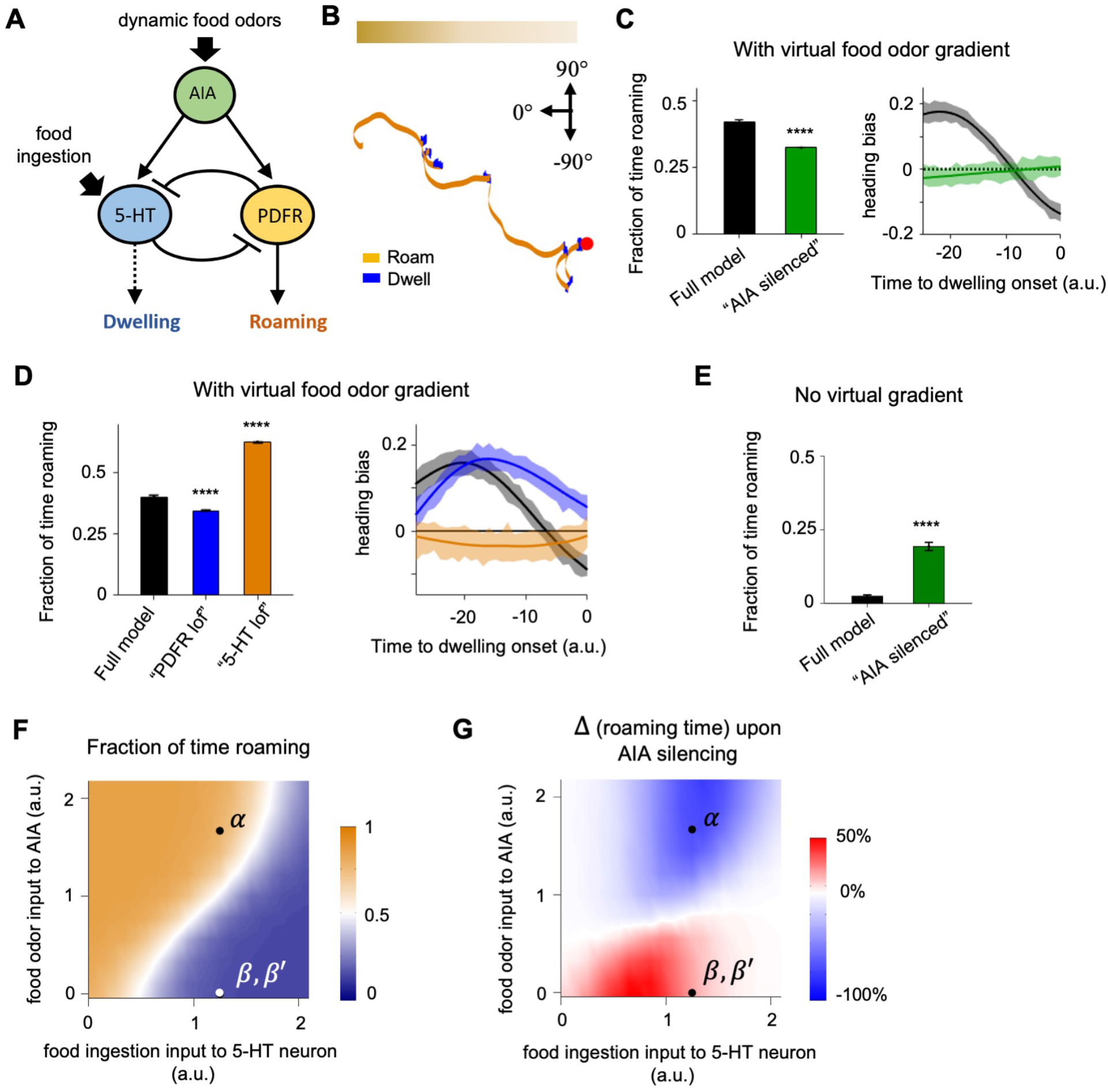
A minimal circuit model suggests that the functional role of AIA depends on combined sensory inputs to the roaming-dwelling circuit. (A) Schematic of the circuit architecture depicted in the model. (B) An example trajectory of an agent migrating up an *in silico* odor gradient (illustrated by the top bar). Red dot indicates the starting location of the animal. The direction pointing up the gradient is designated as 0°. (C) Left: Average fraction of time spent roaming in agents navigating a virtual odor gradient. Agents were driven by either the full circuit model (black) or one where the activity of the model AIA neuron was silenced. Right: Average heading bias (defined the same way as in Fig. 6C) in a time window leading up to the transition from roaming to dwelling. The black curve represents behavior from agents driven by the full circuit model, while the green curve corresponds to the model where the activity of AIA was set to zero. (D) Left: Average fraction of time spent roaming in agents navigating a virtual odor gradient. Agents were driven by either the full circuit model (black) or ones where the activity of the model PDFR neuron (blue) or 5-HT neuron (orange) were attenuated. Right: Average heading bias prior to the transition from roaming to dwelling in agents driven by the full model (black) or ones with the output of the model PDFR neuron (blue) or 5-HT neuron (orange) attenuated. (E) Average fraction of time spent roaming in agents moving in the absence of an odor gradient. Agents were driven by either the full circuit model (gray) or one where the “AIA” neuron is silenced (light green). (F) Average fraction of time spent in the roaming state for agents exposed to various levels of olfactory inputs and ingestion cues. Heat map and color bar indicate the fraction of time spent in the roaming state. Black dots indicate input values taken when the agent moved directly up the gradient (*α*), or in an direction orthogonal or counter to the gradient (*β*). For agents moving in a uniform environment (*β*′), the inputs to the circuit were set to be the same as (*β*). (G) Changes in the fraction of time roaming upon AIA silencing, computed using models where direct sensory input to the model AIA or to the model 5-HT neuron were independently varied. Heat map and color bar indicate increase (red) or decrease (blue) in time spent roaming of the AIA silenced model compared to the full circuit model. Black dots are located in same positions as they are in (F). All error bars are 95% CI. ****p<0.0001, Wilcoxon rank sum test. See Supplemental Methods for additional modeling details.

We simulated the patch foraging assay by providing tonic levels of food ingestion and an olfactory gradient to the simulated agent. Like actual animals, the agents alternated between “roaming” and “dwelling” states as they navigated up the gradient (Fig. 7B). They exhibited elevated levels of roaming compared to no-gradient controls and transitioned into dwelling preferentially when their movement direction started to deviate off the gradient direction (Fig. 7C, compare to Fig. 6B, 6D). When the “AIA” neuron was silenced, the agents exhibited reduced levels of roaming and did not couple their transition into dwelling with their movement direction (Fig. 7C, compare to Fig. 6E). Attenuating the activity of the “5-HT” or “PDFR” neurons resulted in changes in foraging behavior similar to those observed in the corresponding mutant animals (Fig. 7D, compare to Fig. 6F). In contrast, when the agent navigated a uniform environment that only had tonic levels of food input, silencing “AIA” led to an increase in roaming levels, in agreement with experimental results (Fig. 7E, compare to Fig. 6G). These results suggest that a minimal model of this circuit can recapitulate the sensory-dependent control of roaming and dwelling states, as well as our experimental observation that AIA can promote either roaming or dwelling, depending on the sensory context.

We then used this model to examine which set of inputs into the circuit favor AIA promoting roaming versus dwelling. We independently varied the strengths of the dynamic olfactory input to AIA and the ingestion-related cues directly sensed by NSM. We then compared the fraction of time spent roaming in agents driven by the full model to ones where “AIA” was silenced (Fig. 7F-G; also see Methods). With the full model, the level of roaming varied continuously with both inputs, suggesting that the strengths of these two inputs together determine the balance between roaming and dwelling (Fig. 7F). The effect of “AIA” silencing also depended on the levels of both inputs (Fig. 7G). In general, “AIA” promoted roaming when the olfactory input was high and promoted dwelling when the olfactory input was low. AIA’s contribution to roaming was evident at higher ingestion levels, while its role in promoting dwelling became prominent at intermediate to low ingestion levels. These results provide a qualitative match to our experimental data, where AIA promoted roaming as animals navigated up food odor gradients, but promoted dwelling in the absence of such gradients. Taken together, these results suggest that this circuit architecture could allow animals to flexibly select behavioral states in a manner that might allow them to adapt to a wide range of different sensory contexts.

## DISCUSSION

Our findings reveal the functional architecture of a neural circuit that generates persistent behavioral states. Circuit-wide calcium imaging during roaming and dwelling identified stable activity patterns that correspond to each state. By combining circuit imaging with genetic perturbations, we found that mutual inhibition between the serotonergic NSM neuron and the 5-HT and PDF target neurons promotes the stability and mutual exclusivity of these two opposing network states. Further, we found that AIA sensory processing neuron that responds to food odors sends parallel outputs to both neuromodulatory systems and biases the network towards different states in different sensory contexts. This circuit architecture allows *C. elegans* to exhibit persistent roaming and dwelling states, while flexibly switching between them depending on the sensory context.

### Neural circuit mechanisms that generate persistent activity states

The circuit architecture uncovered here provides new insights into how circuits generate persistent activity patterns. Previous work had shown that 5-HT and PDF were critical for dwelling and roaming behaviors^28^, but how they impact circuit activity was not known. We found that NSM and AVB, which produce 5-HT and PDF-1 respectively, have mutually-exclusive activities that correlate with dwelling or roaming. Genetic analyses revealed that the neuromodulators themselves underlie these winner-take-all circuit dynamics. *tph-1* mutants that lack 5-HT had an imbalance in the winner-take-all dynamics, such that NSM activity was less persistent. *pdfr-1* mutants that lack PDF signaling displayed ectopic co-activation of NSM neurons along with AVB and other roaming-active neurons, as well as exaggerated persistence in NSM. These results suggest that neuromodulation is critical to establish the overall structure of circuit-level activity. Our data also suggest that there is mutual inhibition between NSM and the neurons that express MOD-1 (an inhibitory 5-HT receptor) and PDFR-1. The MOD-1- and PDFR-1-expressing neurons, which are active during roaming, synapse onto the PDF-producing neuron AVB that is also active during roaming, suggesting that they excite AVB. Thus, although NSM and AVB display mutually exclusive activities and produce opposing neuromodulators, they do not have direct connections with one another, as is typical in a flip-flop switch. Instead, they coordinate their activities by both interacting with the same network of neurons that expresses the 5-HT and PDF receptors. This architecture might allow for more flexible regulation of behavioral state switching.

The circuit states that correspond to roaming and dwelling differ in several respects. Dwelling states are characterized by persistent activity in serotonergic NSM neurons and reduced activity in several, but not all, locomotion-associated neurons. Roaming states are characterized by fast fluctuations in the activities of neurons that drive forward (AVB, AIY, RIB) and reverse (AVA) movement. The neural dynamics described here, captured by PCA, are distinct from those described in previous studies of non-feeding and immobilized *C. elegans*^5,11,32,33^. PC1 in our data, which correlates with forward-reverse movement, is similar to PC1 from previous studies^5,11^. However, PC2, which correlates with dwelling, was not observed at all in previous studies. This suggests that the *C. elegans* neural activity manifold is reliably constrained in some respects, but also varies considerably across environmental conditions. We did not identify a neuron that is persistently active throughout roaming in a manner analogous to NSM activation during dwelling. While it is possible that such a neuron may exist (and that we did not record it in our study), it is also possible that the roaming state might be the “default” state of the *C. elegans* network and thus does not require devoted, persistently-active neurons to specify the state. Consistent with this possibility, circuit dynamics similar to roaming are observed in the absence of food and even in immobilized animals^11,32,33^. The correlational structure of neural activity also differs between roaming and dwelling. For example, the sensory processing neuron AIA is active in both states, but is coupled to NSM during dwelling, and to the forward-active neurons (AVB, AIY, RIB) during roaming. Neurons that can affiliate to different networks and switch their affiliations over time have also been observed in the stomatogastric ganglion and other systems^57^. The correlational changes that we observe here might allow for state-dependent sensory processing.

### Sensory control of roaming and dwelling states

Previous work showed that chemosensory neurons regulate roaming and dwelling behaviors: mutants that are broadly defective in chemosensation display excessive dwelling, while mutants that are defective in olfactory adaptation display excessive roaming^24^. However, the neural circuitry linking sensory neurons to roaming and dwelling had not been characterized. Using a machine learning-based approach, we identified AIA as a pivotal neuron for roaming-dwelling control. AIA receives synaptic inputs from almost all chemosensory neurons in the *C. elegans* connectome and displays robust responses to appetitive food odors^53,55^. Here we found that AIA provides dual outputs to both the dwelling-active NSM neuron and the roaming-active neurons. Three lines of evidence support this interpretation: (1) native AIA activity correlates with NSM during dwelling and with forward-active neurons during roaming, (2) optogenetic activation of AIA can drive behaviors typical of both states, and (3) AIA silencing strongly alters roaming/dwelling states, but has different effects in different sensory contexts: AIA is necessary for roaming while animals navigate up food odor gradients, but is necessary for dwelling while animals are in uniform feeding environments. Thus, AIA is required to couple the sensory environment to roaming and dwelling states.

The dual output of AIA onto both roaming and dwelling circuits is an unusual aspect of the circuit architecture uncovered here. However, similar functional architectures, where a common input drives competing circuit modules, have been suggested to underlie behavior selection in other nervous systems^1,22,58^. One possible function of this motif in the roaming-dwelling circuit is that it could allow both the roaming- and dwelling-active neurons to be latently activated when the animal is exposed to food odors detected by AIA. AIA-transmitted information about food odors could then be contextualized by other sensory cues that feed into this circuit. For example, NSM is not directly activated by food odors, but instead is directly activated by the ingestion of bacteria via its sensory dendrite in the alimentary canal^39^. Thus, when animals detect an increase in food odors that is accompanied by increased ingestion, this might promote dual AIA and NSM activation to drive robust dwelling states. In contrast, when animals detect an increase in food odors that is not accompanied by increased ingestion, this might activate AIA and the other side of the mutual inhibitory loop, biasing the animal towards roaming. This flexible architecture could therefore allow animals to make adaptive foraging decisions that reflect their integrated detection of food odors, food ingestion, and potentially other salient sensory cues.

## MATERIALS AND METHODS

### Growth conditions and handling

Nematode culture was conducted using standard methods^59^. Populations were maintained on NGM agar plates with *E. coli* OP50 bacteria. Wild-type was *C. elegans* Bristol strain N2. For genetic crosses, all genotypes were confirmed using PCR. Transgenic animals were generated by injecting DNA clones plus fluorescent co-injection marker into gonads of young adult hermaphrodites. One day old hermaphrodites were used for all assays. All assays were conducted at room temperature (∼22°C).

### Strain List

| Genotype | Source | Identifier |
| --- | --- | --- |
| Wild-type Bristol N2 | N/A | N2 |
| *flvEx46*[*tph-1::GCaMP6m, mod-1::GCaMP6m, sto-3::GCaMP6m, glr-3::GCaMP6m, odr-2b::GCaMP6m, gcy-28.d::GCaMP6m, lgc-55(short)::GCaMP6m, nmr-1::GCaMP6m, tph-1::wormScarlett, mod-1::wormScarlett, nmr-1::wormScarlett, sto-3::wormScarlett*]; *lite-1(ce314), gur-3(ok2245)*] | This study | SWF90 |
| *flvIs1*[*tph-1::GCaMP6m, mod-1::GCaMP6m, sto-3::GCaMP6m, glr-3::GCaMP6m, odr-2b::GCaMP6m, gcy-28.d::GCaMP6m, lgc-55(short)::GCaMP6m, nmr-1::GCaMP6m, tph-1::wormScarlett, mod-1::wormScarlett, nmr-1::wormScarlett, sto-3::wormScarlett*]; *lite-1(ce314), gur-3(ok2245)*] | This study | SWF113 |
| *flvIs1; lite-1(ce314); gur-3 (ok2245); mod-1(ok103)* | This study | SWF186 |
| *flvIs1; lite-1(ce314); gur-3 (ok2245); pdf-1(ok3425)* | This study | SWF124 |
| *flvIs1; lite-1(ce314); gur-3 (ok2245); flvEx129*[*pdf-1::acy-1gf, elt-2::nGFP*] | This study | SWF263 |
| *flvEx46; lite-1(ce314); gur-3 (ok2245); tph-1(mg280); pdf-1(ok3425)* | This study | SWF125 |
| *flvIs1; lite-1(ce314); gur-3 (ok2245); flvEx103*[*tph-1(short)::Chrimson, elt-2::nGFP*] | This study | SWF197 |
| *kyls580*[*mod-1::nCre, myo-2::mCherry*]; *kyEx4816*[*ttx-3::ChR2(C128S)-GFP, odr-2b::inv[ChR2-sl2-GFP], myo-3::mCherry*] | This study | CX14686 |
| *pdf-1(ok3425); kyls580*[*mod-1::nCre, myo-2::mCherry*]; *kyEx4816*[*ttx-3::ChR2(C128S)-GFP, odr-2b::inv[ChR2-sl2-GFP], myo-3::mCherry*] | This study | CX14684 |
| *flvIs1; lite-1(ce314); gur-3 (ok2245); flvEx98*[*gcy-28.d::Chrimson, elt-2::nGFP*] | This study | SWF194 |
| *flvIs1; lite-1(ce314); gur-3 (ok2245); tph-1(mg280); flvEx98*[*gcy-28.d::Chrimson, elt-2::nGFP*] | This study | SWF216 |
| *flvIs1; lite-1(ce314); gur-3 (ok2245); pdf-1(ok3425); flvEx98*[*gcy-28.d::Chrimson, elt-2::nGFP*] | This study | SWF326 |
| *kyEx4745*[*gcy-28.d::unc-103gf::sl2-mCherry, elt-2::mCherry*] | Larsch et al., 2015 | CX14597 |
| *tax-4(p678) [5x backcrossed to N2]* | This study | CX13078 |
| *pdf-1(ok3425)* | Flavell et al., 2013 | CX14295 |
| *tph-1(mg280)* | CGC | MT15434 |
| *flvEx54*[*gpa-6::CoChR, myo-3::mCherry*] | This study | SWF103 |
| *flvEx54*[*gpa-6::CoChR, myo-3::mCherry*]; *kyEx4745*[*gcy-28.d::unc-103(gf)::sl2-mCherry, elt-2::mCherry*] | This study | SWF315 |

### Construction and characterization of multi-neuron GCaMP strain

To generate a transgenic strain expressing GCaMP6m in a specific subset of neurons involved in roaming and dwelling, we first generated pilot strains where one or two plasmids were injected at a time to optimized DNA concentrations. This also allowed us to determine the precise GCaMP6m and/or Scarlett expression pattern from each promoter. We then injected these plasmids as a mixture into *lite-1;gur-3* double mutants, which are resistant to blue light delivered during calcium imaging. We selected a line for use that had normal behavioral parameters and showed relatively balanced expression of GCaMP6m and Scarlett in the target cells (SWF90). To obtain more consistent expression, the transgene was integrated by UV to generate *flvIs1* (SWF113). The integrated strain was outcrossed 4 times.

### Microscope Design and Assembly

#### Overview

The tracking microscope design was inspired and based off previously described systems^31–33^, with several modifications aimed at reducing motion artifacts and extending the duration of calcium imaging, so that long-lasting behavioral states could be examined. As illustrated in Figure 1-Figure Supplement 1, two separate light paths, below and above the specimen, were built onto a Ti-E inverted microscope (Nikon).

#### High-magnification light path for GCaMP imaging

The light path used to image GCaMP6m and Scarlett at single cell resolution is an Andor spinning disk confocal system. Light supplied from a 150mW 488nm laser and a 50mW 560nm laser passes through a 5000rpm Yokogawa CSU-X1 spinning disk unit with a Borealis upgrade (with a dual-camera configuration). A 40x/1.15NA CFI Apo LWD Lambda water immersion objective (Nikon) with a P-726 PIFOC objective piezo (PI) was used to image the volume of the worm’s head. A custom quad dichroic mirror directed light emitted from the specimen to two separate Andor Zyla 4.2 USB3 cameras, which had in-line emission filters (525/50, and 625/90). Data was collected at 2×x2 binning in a 512×512 region of interest in the center of the field of view.

#### Low-magnification light path for closed-loop tracking

A second light path positioned above the animal collected data for closed-loop tracking. Light supplied from a Sola SE2 365 Light Engine (Lumencor) passed through a DSRed (49005, Chroma) filter set and a 10x/0.3NA air objective to excite Scarlett in the head of the worm. Red light emitted from the specimen passed through the filter set to an acA2000-340km Basler CMOS camera. Data was collected at 100 Hz.

#### Synchronized control of camera exposures and illumination light sources

The Andor Zyla cameras used for calcium imaging were run in rolling shutter mode. A trigger signal was generated by one of the two cameras whenever the camera shutter is fully open (∼2 ms per exposure). This trigger signal served as a master control that synchronized several devices (Figure 1-Figure Supplement 1B). First, it was used to drive the 488nm and 560nm lasers, such that illumination is only provided when the full field of view is open. Second, the same trigger signal was used controlled the movement of the objective piezo, such that fast piezo movement occurs largely outside the window of laser illumination. Lastly, this signal was used to time the green LED used by the closed-loop tracking system. The LED was turned on only when the calcium imaging cameras were not actively acquiring images (i.e. outside the window when the rolling shutter is fully open) and when the lasers were off. Together, these approaches minimize photo-bleaching, photo-toxicity, and motion artifacts induced by movable parts of the microscope.

#### Closed-loop tracking software

A custom C/C++ software was used to process incoming frames from the tracking camera and to instruct the movement of a motorized stage (96S107-N3-LE2, Ludl; with a MAC6000 controller) to keep the head region of the animal at the center in the field of view. This software was adapted from Nguyen et al. with two key modifications: First, at each control cycle, the future velocity of the stage was calculated to match the predicted future velocity of the animal (i.e. predictive control as opposed to proportional control employed in previous study). Specifically,

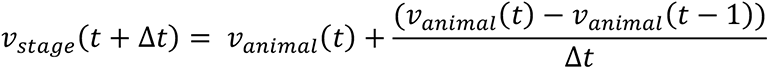

where *v_stage_*(*t*) is the instantaneous velocity of the stage and *v_animal_*(*t*) the instantaneous velocity of the animal. The latter was estimated as described below (see **Estimation of instantaneous animal location and velocity**). The right side of the formula was found empirically to be sufficient for predicting future animal velocity. The second modification was that we used the motion of the head region of the animal to extrapolate the locomotory state of the animal. This approach was empirically justified in a recent study and circumvents the need for a third light path for imaging the full body of the animal.

### Behavioral Assays

#### Patch foraging assay

For the patch foraging assays, we used 24.5cm x 24.5cm NGM plates. Plates were uniformly seeded with sparse OP50 bacteria (OD 0.5 diluted 300x), and one half of the plate was seeded with dense bacteria (OD 0.5 concentrated 20x). The border between the sparse and dense food was always sharp and typically very straight. Plates were left overnight at room temperature. The following day, one-day old adult animals were picked to the sparse side of the food plate, approximately 1.5 cm from the dense food patch. Video recordings were started immediately, though for all analyses the first 20 min of data (equilibration time) was not analyzed. Videos were recorded at 3 fps using Streampix 7.0, a JAI SP-20000M-USB3 CMOS camera (41mm, 5120×3840, Mono) and a Nikon Micro-NIKKOR 55mm f/2.8 lens. Backlighting was achieved using a white panel LED (Metaphase Technologies Inc. White Metastandard 10” X 25”, 24VDC). Assay plates were placed on glass 3” above LEDs to avoid heat transfer to plates. Videos were processed using custom Matlab scripts, which included a step to manually confirm the exact frame of lawn encounter for each animal. Segmentation of behavior into roaming and dwelling was conducted as previously described.

#### Foraging at different food densities

To examine animal behavior in uniform environments with different food densities, we seeded NGM plates (either circular 10 cm or 24.5×24.5cm) with different densities of food. For the experiments in Fig. 6, low-density was OP50 bacteria at OD 0.5 diluted 300x; high-density was OD 0.5 concentrated 20X. Plates grew overnight at room temperature. The following day, one day-old adult animals were picked to these plates and allowed to equilibrate for 45 mins, after which video recordings began. Videos were recorded and analyzed as described above.

#### Optogenetic stimulation during foraging behavior

For optogenetic stimulation of free-behaving animals, we picked one day-old adult animals to NGM plates seeded with 300X diluted OD 0.5 OP50 (supplemented with 50 uM all-trans-retinal) the night before. Animals were permitted to equilibrate for 45 min, after which videos were recorded using the setup described above. In these videos, light for optogenetic stimulation was delivered using a 625nm Mightex BioLED at 30 uW/mm^2^. Patterned light illumination was achieved using custom Matlab scripts, which were coupled to a DAQ board (USB-6001, National Instruments) and BioLED Light Source Control Module (Mightex). Videos were analyzed as described above.

### Data Analysis for Calcium Imaging

#### Semi-automated image segmentation to obtain neuron outlines

All image analyses were performed on maximum intensity projections of the collected z-stacks, since the neurons were well separated along the x-y axes. First, feature points and feature point descriptors were extracted for each frame of the calcium imaging video. Next, an N-by-N similarity matrix (N = number of frames in a video) was generated where each entry equals the number of matched feature points between a pair of frames. The columns of matrix were clustered using hierarchical clustering. Around 30 frames (typically 1-2% of frames from a video) were chosen across the largest 15 clusters. These frames were then segmented manually. The user was asked to outline the region for interest (ROI) around each neuronal structure of interest (axonal segment for the AIY neurons, soma for all other neurons). After manual segmentation, the automatic segmentation software loops through each of the remaining frames. For each unsegmented frame (target frame), a best match (reference frame) was found among the segmented frames based on the similarity matrix. Then, geometrical transformation matrices were estimated using the locations of the matched feature points. The estimated transformation was then applied to the boundary vertices of each ROI in the reference frame to yield the estimated boundary of the same region in the target frame. Once done, the target frames with its automatically computed ROIs was included into the pool of segmented frames and could serve as a reference frame for the remaining unsegmented frames. This procedure was repeated iteratively through the rest of the video.

#### Estimation of instantaneous animal location and velocity

The instantaneous location of the animal 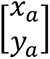 was calculated based on the following formula:

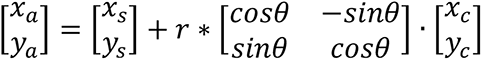

where 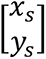 is the instantaneous location of the microscope stage, 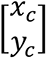 is the position of the head region of the animal as seen on the frame captured by the tracking camera, *θ* is the rotation angle between the field of view of the tracking microscope and the sensor of the tracking camera, and *r* is the pixel size of the frames taken through the tracking camera. The velocity of the animal calculated by dividing the displacement vector of the animal between adjacent time points by the duration of the time interval.

#### Aligning calcium imaging data with behavioral data

As described in the Microscope Design and Assembly section, the trigger signal for the confocal laser was simultaneously sent to the computer controlling the tracking microscope. This computer thereby store two sets of time stamps, one for the laser illumination sequence and the other for the behavioral tracking video. Since the internal clock is the same, we can interpolate both the calcium activity data and the behavioral data onto the same time axis. Specifically, we interpolated both the calcium activity and behavior time series to obtain a common sampling frequency of 2 Hz.

#### Calcium signal extraction and pre-processing

The fluorescent signal from each neuron in a given frame was calculated as the median of the brightest 100 pixels within the ROI (or all pixels if the size of the ROI was smaller than 100 pixels) of that neuron. This approach was adopted to render the estimation of calcium signal insensitive to the exact segmentation boundary of the neuron, which could inadvertently contain background pixels. This was done for both the green and the red channels. The following pre-processing steps were then applied to the time-series signals from both channels: 1) To reduce spurious noise, a sliding median filter with a window size of 5 frames were applied to the time series (Figure 1-Figure Supplement 2D). 2) To correct for the decay in fluorescent signal due to photobleaching, an exponential function was first fit to the time series. Next, the estimated exponential was normalized by its initial value and divided away from the denoised time series (Figure 1-Figure Supplement 2E). 3) To control for fluctuations in fluorescent signal due to the movement of the animal, we calculated the ratiometric signal. Specifically, the denoised and bleach-corrected time series from the green channel was divided by that from the red channel. 4) Lastly, to control for the variations in the dynamic range of the calcium signal due to variations in the expression of the fluorescent indicators, we normalized the ratio-metric signal such that the 1^st^ percentile of the signal takes a value of 0 while the 99^th^ percentile takes the value of 1. To control for cases where a given neuron never became active in a given recording (e.g. NSM in *pdfr-1::acy-1gf* animals), exceptions were made if a neuron’s peak activity in a given recording was less than 10% of the average across all recordings. In this case, the original ΔR/R0 value was used without normalization. Apart from this exception, the normalized ratio-metric signal was used for all subsequent data analyses. These data processing steps (dividing by mScarlett; normalizing to a 0-1 scale) did not change the distributions of GCaMP intensity values (Figure 1-Figure Supplement 2H).

#### Principal component analysis (PCA)

An *N-by-M* data matrix was assembled with the rows representing neuron identity (*N* = 10) and the columns time points. For each wild-type data set, the calcium activity data from each neurons was represented as row vectors and concatenated along the neuron dimension. Data across different recording sessions were concatenated along the time dimension. PCA was performed by first subtracting the mean from each row and then applying singular value decomposition to the matrix. We chose this method over the previously described approach of performing PCA on the time derivatives of the calcium signals^11^. This is because we found that applying PCA on the time derivatives did not yield PCs with intuitive behavioral correlates when applied to our data (Figure 1-Figure Supplement 3C), potentially because the timescales of the neural dynamics were notably different in foraging animals, versus previous studies.

#### Cross-correlation in neural activity

To estimate the time-lagged similarity between the activity of two neurons for a given genotype, the cross-correlation function (XCF) was first calculated individually for each data set of that genotype and then averaged. Bootstrapping was done to obtain confidence intervals on the mean. To examine the functional coupling between two neurons over time, average XCFs were calculated for data from a series of 60 second time windows spanning from 90 seconds before NSM activation to 90 seconds after. For each time window, the point with the largest absolute value along the average XCF was identified. The mean and 95% CI values of these extrema points were concatenated chronologically to generate plots.

#### Convolutional neural network (CNN) classifier

The classifier was implemented using the Deep Learning Toolbox in MATLAB. The architecture of the network consists of a single convolutional layer with a single channel of two 9-by-3 convolutional kernels with no padding, followed by a Rectified Linear Unit (ReLu) layer, a fully connected layer with two neurons, a two-way softmax layer and a classification output layer. The last layer is specifically required for the Matlab implementation and computes the cross-entropy loss. Calcium activity from all neurons imaged, except for the 5-HT neuron NSM, were used for training, validation and testing. To specifically predict transition from roaming to dwelling, only data during roaming were used to predict the onset of NSM activity. For each wild-type data set, calcium activity during each roaming state was first down-sampled by applying a 30 second average filter starting from right before the onset of a dwelling state and going back in time to the beginning of the roaming state. Each time point in the down-sampled data was assigned a label of 1 or 0: 1 if it is immediately prior to an episode of NSM activation, and 0 otherwise. Positive and negative samples were balanced by weighting the prediction error of each sample by the number of samples in that class. The positive and negative sample groups were each partitioned at random into the training, validation, and test sets at an 8:1:1 relative ratio. This random partition was repeated 200 times. For each data partition, network training was performed 10 times with random initial conditions, using Stochastic Gradient Descent with Moment (SGDM) with the following hyper-parameters:

| Hyper-parameter name | Value |
| --- | --- |
| Initial Learning Rate | 0.09 |
| L2 Regularization Rate | 0.0001 |
| Learning Rate Drop Factor | 0.1 |
| Learning Rate Drop Period | 10 |
| Momentum | 0.9 |
| Validation Frequency | 30 |
| Max number of epochs | 150 |

To identify convolutional kernels that consistently contribute to classifier accuracy, convolutional kernels from networks that achieved greater than 50% test accuracy were recorded and k-means clustering was performed. Within each cluster, the distribution of weights at each kernel location was used to extract a confidence interval for the mean value of that kernel element. Elements of the kernel with mean values significantly different from 0 were taken to indicate important neural activity profiles for predicting NSM activation. Since each kernel element maps to the activity of a given neuron at a particular time window, the preferred sign of a kernel element would suggest whether a neuron is preferentially active (when the preferred sign is positive) or inactive (when the preferred sign is negative) at that time window.

Feature selection was performed to identify key neurons whose activity critically contribute to classification accuracy. To generate the results in Fig. 4B, data from a chosen neuron was removed from the 9-neuron data set, and the resulting partial data set was used to train CNNs following procedure described above. To generate the results in Figure 4-Figure Supplement 1B, two types of partial data sets were used. In the first category, data from 6 out of 9 neurons were used for training. We tested all possible 9-choose-6 neuron combinations. In second category, we tested using data from only RIB, AIA, and AVA for network training.

### Data Analysis for Behavioral Assays

#### Extraction of locomotory parameters

Animal trajectories were first extracted using custom software described previously ^39^. Speed and angular speed were calculated for all time points of each trajectory, and then averaged over 10 second intervals.

#### Identification of roaming and dwelling states

Roaming and dwelling states were identified as previously described ^28^. Briefly, the speed and angular speed measured for each animal at each time point was assigned into one of two clusters. This allowed each animal trajectory to be converted into a binary sequence. A two-state HMM was fit to these binary sequences to estimate the transition and emission probabilities. This was done separately for each genotype under each experimental condition.

#### Calculation of heading bias

The instantaneous heading bias *c*(*t*) was defined as:

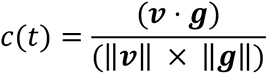

Where *v* is the instantaneous velocity of the animal, and *g* is the unit vector that points from the animal’s current location to the nearest point on the boundary between the sparse food patch and the dense food patch. Here, *g* is used as the proxy for the gradient of olfactory cues at the animal’s current location. Equivalently, *c*(*t*) is the cosine of the angle between the animal’s instantaneous direction of motion and the direction of the chemotactic gradient at its current location.

### Modeling

#### Neural circuit model

##### Model setup

To examine the general properties of the 3-node circuit motif shown in Fig. 7A, we constructed the following model:

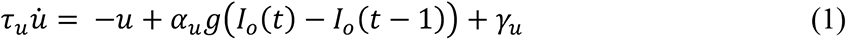

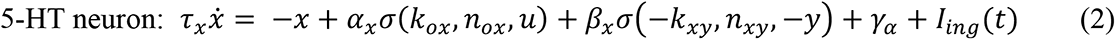

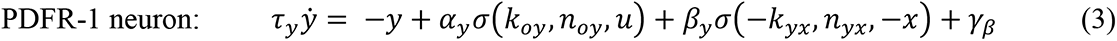

where *g*(*x*) = *xH*(*x*) and 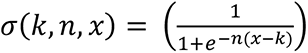, with (*x*) being the Heaviside function. *I_o_*(*t*) is the concentration of food-odor cues experienced by the animal at time *t*, while *I_ing_*(*t*) is the ingestion-related cues sensed directly by the 5-HT neuron NSM. This formulation of AIA dynamics is motivated by recent studies, which showed that AIA activity appears as a low-pass filtered version of the temporal difference in odor concentration experienced by the animal^60,61^.

##### Model reduction

Because AIA activity peaks rapidly upon activation and its peak amplitude reliably scales with the temporal difference in odor concentration^60,61^, we treat AIA activity as a proxy for the gradient-related sensory input and approximate (1) with its steady state solution:

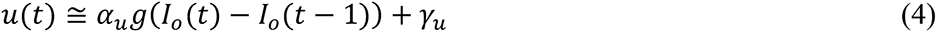

We further define *I_g_*(*t*) = *g*(*I_o_*(*t*) − *I_o_*(*t* − 1)) as the temporal difference of olfactory inputs and substitute (4) into (2) and (3). This reduces the circuit model to:

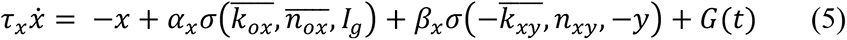

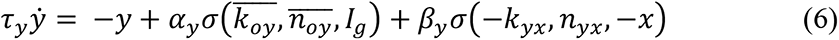

##### Simulation of foraging behavior

To simulate the patch foraging assay, agents were exposed to a linear olfactory gradient that lies parallel to the x-axis: *I_o_*(*t*) = *kx*(*t*). To simulate the control assay with a uniform environment, we set *I_o_*(*t*) to be constant everywhere.

The foraging behavior of individual agents was generated using the following pseudocode:

0. At *t*_0_, the agent is in the roaming state. It is located at the starting location(0,0). The initial condition of the neural circuit model was set to be at the steady-state values when the PDFR-1 neuron dominates (the “PDFR high” state).
1. At *t_n_*(*n* ≥ 1), the agent moves at a fixed roaming speed *v_r_* for a fixed duration *dt*. The heading angle *θ*_0_ is drawn randomly from a uniform distribution *U*(−*π*, *π*). The time step size is *dt*.
2. The model then computes the change in the level of olfactory cues between the agent’s current and prior location at *t_n-1_*. The resulting value serves as the gradient-related olfactory input (*I_g_*) to the neural circuit model.
3. The activity of the 5-HT and PDFR-1 neurons are computed according to eqs. 5 and 6 using the Runge-Kutta (RK4) method.
4. The updated circuit activity is compared against the pre-computed separatrix of the dynamical system described by eqs. 5 and 6. Depending on which side of the separatrix the circuit activity values lie, the circuit is designated to be in a “PDFR high” or a “5-HT high” state. The state of the circuit in turn determines whether the agent will be in the roaming or dwelling states, respectively, in the next time step (*t_n+1_*).
5. At the beginning of a roaming state, the instantaneous heading angle of the agent is chosen at random from *U*(−*π*, *π*). From then on, the heading angle evolves according to *θ*(*t*) = *sgn*(*θ*(*t* − 1)) ∗ (|*θ*(*t* − 1)| − φ + *η*_*θ*_(*t*)). The noise term *η*_*θ*_(*t*) is a noise term drawn from a normal distribution Ν(0, ε).
6. Repeat step 1-4 until the end of simulation *t_N_*.

##### Parameter selection

To explore the general properties of the circuit architecture shown in Fig. 7A, we performed a parameter screen to identify parameter regimes where AIA-mediated olfactory input greatly enhances the probability of entry into the dwelling state. We sample the following parameters were independently sampled from uniform distributions described below:

| Parameter | Sampling distribution | Notes |
| --- | --- | --- |
| $\alpha_x$ | $U(0,5)$ | This sets the max increase in activity from gradient-related olfactory input |
| $\alpha_y$ | $U(0,5)$ | Ibid. |
| $\beta_x$ | $U(1,5)$ | This sets the difference in activity between the fully inhibited and the fully disinhibited states of the neuron |
| $\beta_y$ | $U(1,5)$ | Ibid. |
| $k_{ox}$ | $U(0,5)$ | This sets the value of olfactory input that evokes half-maximum excitation |
| $k_{oy}$ | $U(0,5)$ | Ibid. |
| $n_{ox}$ | $U(0,6)$ | This sets the steepness of the sigmoid curve. $n = 6$ correspond to a near step-function-like curve |
| $n_{oy}$ | $U(0,6)$ | Ibid. |
| $k_{xy}$ | $U(0,5)$ | This sets the value of $y$ that evokes half-maximum inhibition on $x$ |
| $k_{yx}$ | $U(0,5)$ | This sets the value of $x$ that evokes half-maximum inhibition on $y$ |
| $n_{xy}$ | $U(1.5,6)$ | This sets the steepness of the sigmoid curve. $n > 1$ is required for bi-stability from mutual inhibition. $n = 6$ correspond to a near step-function-like curve |
| $n_{yx}$ | $U(1.5,6)$ | Ibid. |

We held the initial condition of the circuit and the inputs to the circuit at constant values. In addition, we had assumed that the 5-HT neuron and PDFR neurons operate at similar time scales:

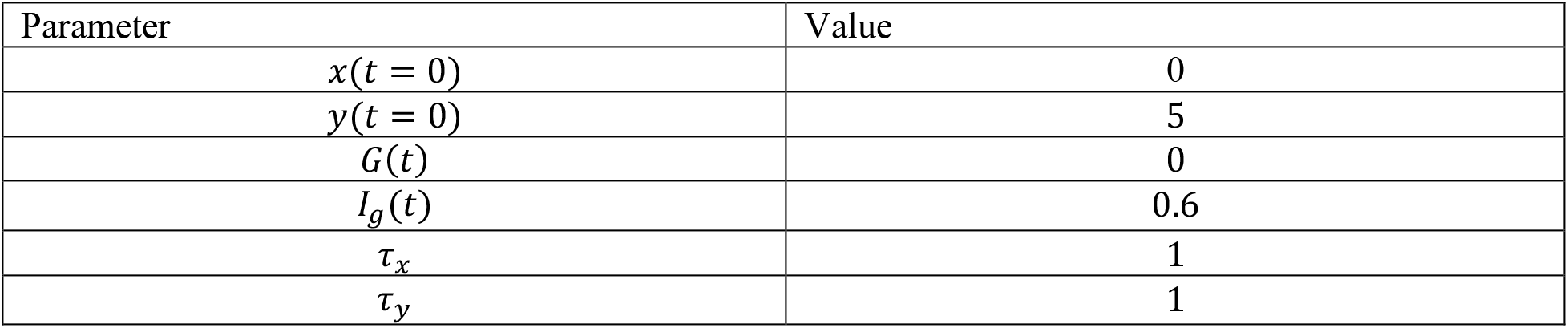

For each set of randomly generated model parameters, we estimate the probability of the circuit entering a “dwelling-like”, or “5-HT high” state by simulating the circuit model (eqs. 5 and 6) with Gaussian noise (*N*(0,0.25)) injected into both circuit nodes. The simulation was kept short (50 time steps) and repeated 200 times to compute the probability that the circuit entered the “5-HT high” at least once during the duration of the simulation. For comparison, we then repeat the simulation with a modified circuit model where the AIA-dependent olfactory input onto the 5-HT neuron is blocked (i.e. setting *I_g_*(*t*) ≡ 0 for eq. 5). We then re-compute the probability of entering “5-HT high” state for the modified model and compare it with the original model. Parameter sets that result in a large increase in the probability of entering the “5-HT high” state were used to drive the simulation of foraging behavior in virtual agents. The following parameter values were used for simulations shown in Fig.7*:

| Parameter | Value |
| --- | --- |
| $\alpha_x$ | 1.6 |
| $\alpha_y$ | 3.6 |
| $\beta_x$ | 4.0 |
| $\beta_y$ | 3.7 |
| $k_{ox}$ | 0.4 |
| $k_{oy}$ | 1.0 |
| $n_{ox}$ | 4 |
| $n_{oy}$ | 5 |
| $k_{xy}$ | 4.2 |
| $k_{yx}$ | 3.2 |
| $n_{xy}$ | 2 |
| $n_{yx}$ | 2 |

### Statistical Analysis

#### Comparison of sample means

The Wilcoxon ranksum test was applied pair-wise to obtain the raw p-values. When multiple comparisons were done for the same type of experiment (e.g. comparing the fraction of animal roaming during the patch foraging assay for different genotypes), the Benjamini-Hochberg correction was used to control the false discovery rate. A corrected p-value less than 0.05 was considered significant.

#### Bootstrap confidence intervals

Bootstrapping was performed by sampling with replacement *N* times from the original data distribution (*N* equals the size of the original distribution). This procedure was repeated 1000 times and the test statistic of interest (e.g. the sample mean) was calculated each time on the bootstrapped data. The 5^th^ and 95^th^ percentiles of the calculated values then constitute the lower and upper bounds of the 95% confidence interval.

## AUTHOR CONTRIBUTIONS

N.J. and S.W.F conceived of the study and wrote the paper. N.J., G.K.M., and S.W.F. designed/performed experiments and performed data analysis. N.J., G.I.F., and A.D. designed and wrote software. C.M.B., and I.N. performed experiments and analyzed data.

## DECLARATION OF INTERESTS

The authors have no competing interests to declare.

## ACKNOWLEDGMENTS

We thank Rachel Wilson, Andrew Gordus, Paul Greer, Yun Zhang, Michael Hendricks, Mike O’Donnell, Dipon Ghosh, and members of the Flavell lab for helpful comments on the manuscript. We thank Andrew Leifer for helpful advice and sharing software related to the tracking microscope, Thomas Boulin for sharing the mScarlett plasmid, and Nate Cermak for help with hardware control on the tracking microscope. We thank the Bargmann lab and the *Caenorhabditis* Genetics Center (supported by P40 OD010440) for strains. N.J. acknowledges support from the Picower Fellows program and the Charles King Trust Postdoctoral Fellowship. S.W.F. acknowledges funding from the JPB Foundation, PIIF, PNDRF, the NARSAD Young Investigator Award Program, McKnight Foundation, NIH (R01NS104892) and NSF (IOS 1845663 and DUE 1734870).

## SUPPLEMENTARY FIGURE LEGENDS

**Figure 1 – Figure Supplement 1.**
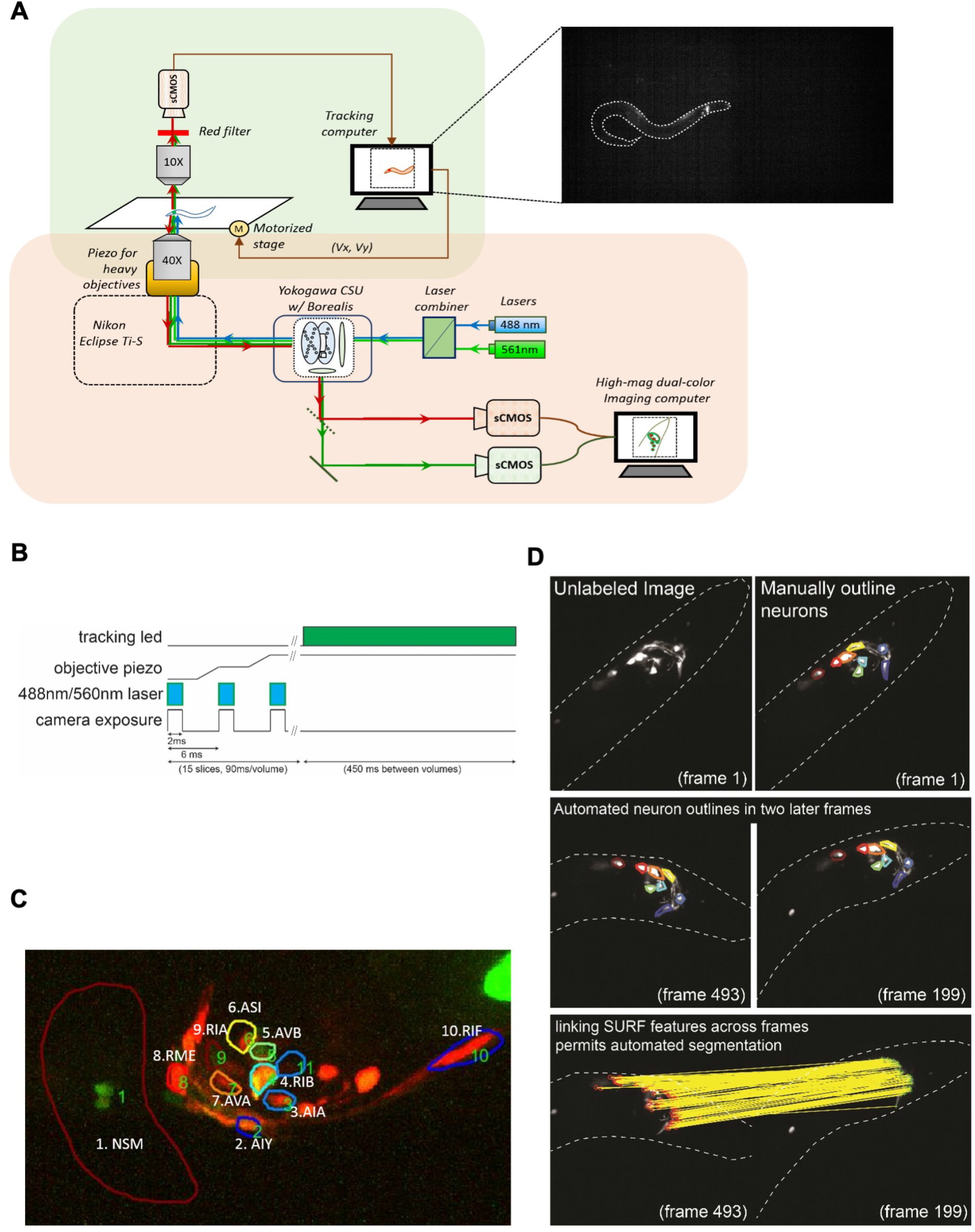
Design and calibration of the spinning-disk confocal tracking scope. (A) Design of the microscope. Orange and green shaded boxes indicate the confocal and behavioral tracking parts of the microscope, respectively. An example image from the behavior tracking camera is shown, with the worm outlined in white. mScarlett-expressing neurons can be robustly detected in the animal’s head. (B) To minimize photo-bleaching, movement artifacts, and animal disturbance, the laser illumination of animals was timed to camera exposure and objective piezo movement during volume acquisition, as is illustrated. The tracking LED was also only illuminated in between GCaMP/mScarlett volume acquisitions, so as to prevent cross-talk between the upper and lower microscope paths. Laser illumination permitted animal tracking during volume acquisition. (C) A sample volume captured by the confocal microscope. Neurons expressing the GCaMP6m and the mScarlett fluorescent proteins are annotated. For AIY, the neurite is labeled. (D) Semi-automated segmentation of neuron boundaries using the SURF algorithm. For a subset of frames in a video, the neuron boundaries are manually outlined. Then, the boundaries are propagated from one frame to others, based on image transformations that are defined by matching SURF features across frames.

**Figure 1 – Figure Supplement 2.**
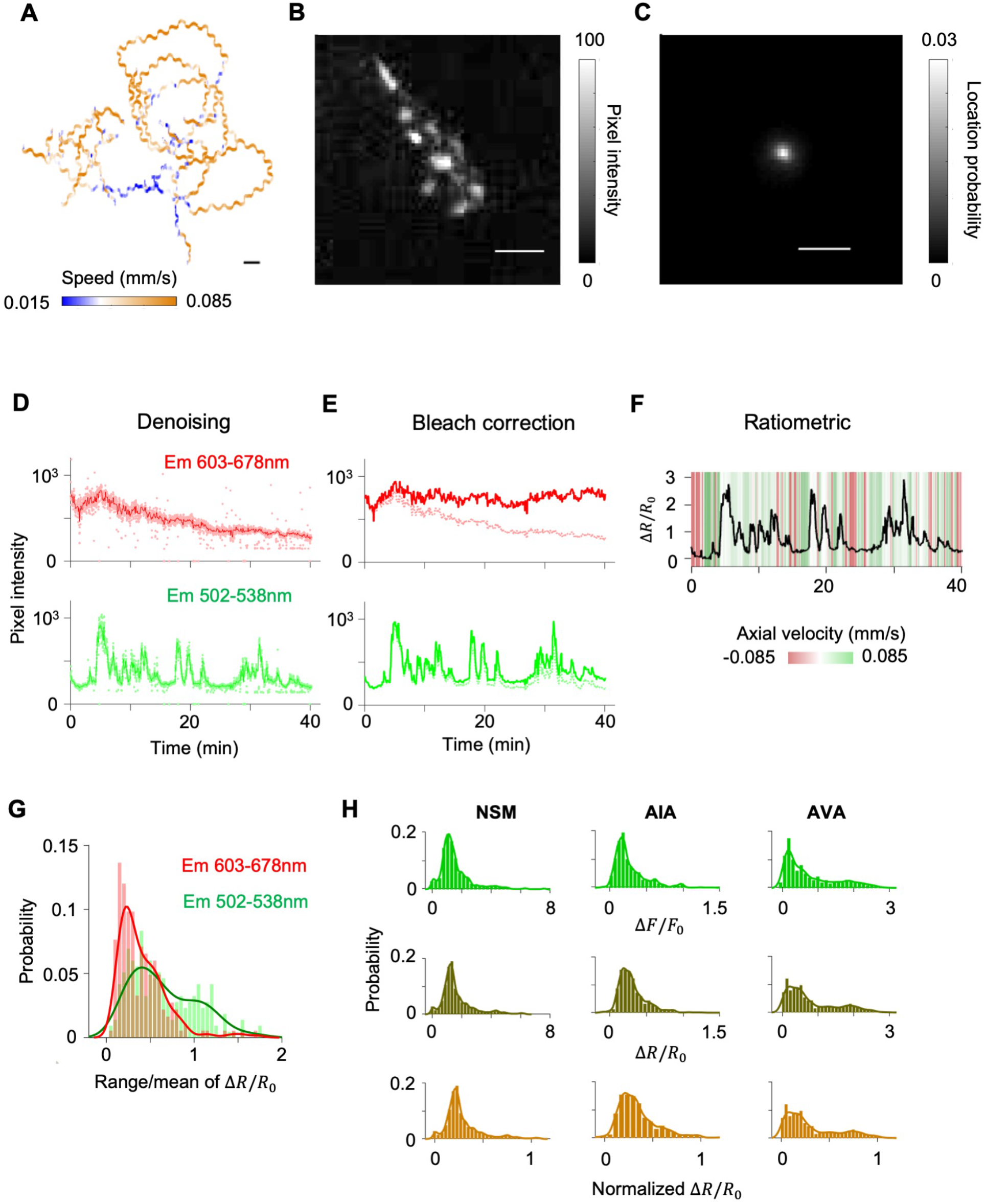
Calibration of behavioral tracking accuracy and the effect of motion on calcium imaging data. (A) Example trajectory of an animal recorded under the tracking confocal microscope. (B) An image of this animal’s head region captured through the behavior tracking camera. Bright pixels correspond to neurons expressing the mScarlett transgene. (C) Probability distribution of the location of the head region as seen through the behavior tracking camera across all of the frames of the recording shown in (A). All scale bars in (A-C) represent 0.1 mm. (D-F) Extraction of calcium activity from dual-channel fluorescent intensities from a representative neuron. (D) Time series of red (emission wavelength 603-678nm) and green (emission wavelength 502-538nm) fluorescent intensities were first denoised by median filtering. (E) Next, photobleaching over time was corrected by fitting and then normalizing away an exponential decay function. (F) Finally, the time series data from the green channels was divided by that from the red channel. The resulting time series was normalized to a relative scale of 0 to 1, with 0 corresponding to the 1st percentile and 1 to the 99th percentile of ratiometric values. (G) Range of variation normalized by mean calculated for the bleach-corrected red and green fluorescent intensities. Histograms were computed for aggregate data from all videos used in this study. Curved lines overlaying the red and green histograms (color matched) are mixture of Gaussian models fit to the corresponding histogram. (H) Distributions of fluorescence measurements are not perturbed by data processing. Probability distribution functions for the activity of three example neurons after various stages of data pre-processing: (top) cell-specific fluorescent signals from the green channel after denoising, bleach correction and baseline subtraction; (middle) ratiometric values computed by dividing signals from the green channel with those from the red channel; (bottom) normalized Δ*R*/*R_o_* values computed by remapping the 1^st^ and 99^th^ percentiles of the distribution to 0 and 1.

**Figure 1 – Figure Supplement 3.**
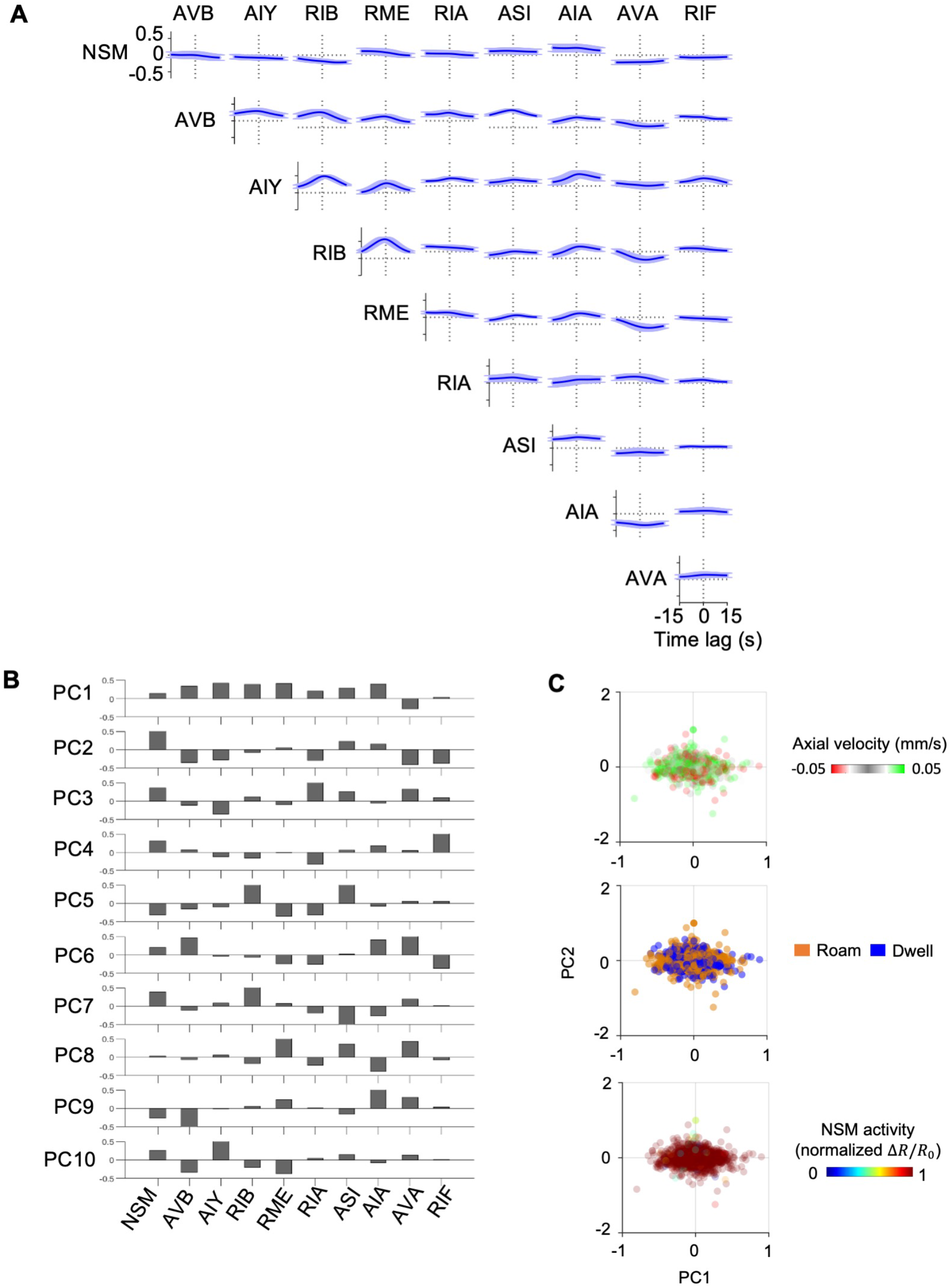
PCA analysis of wild-type circuit activity. (A) Average pair-wise cross-correlation function among neurons examined through free-moving calcium imaging of wild-type animals (N=17). Data are shown as means and standard error. (B) Loadings of individual neurons on to PCs 1-10. (C) PCA calculated using the temporal derivative of the neural activity data. Note that data do not segregate based on behavior when performing PCA in this manner. Color scale represents (top) axial velocity, (middle) foraging state, or (bottom) NSM activity.

**Figure 1 – Figure Supplement 4.**
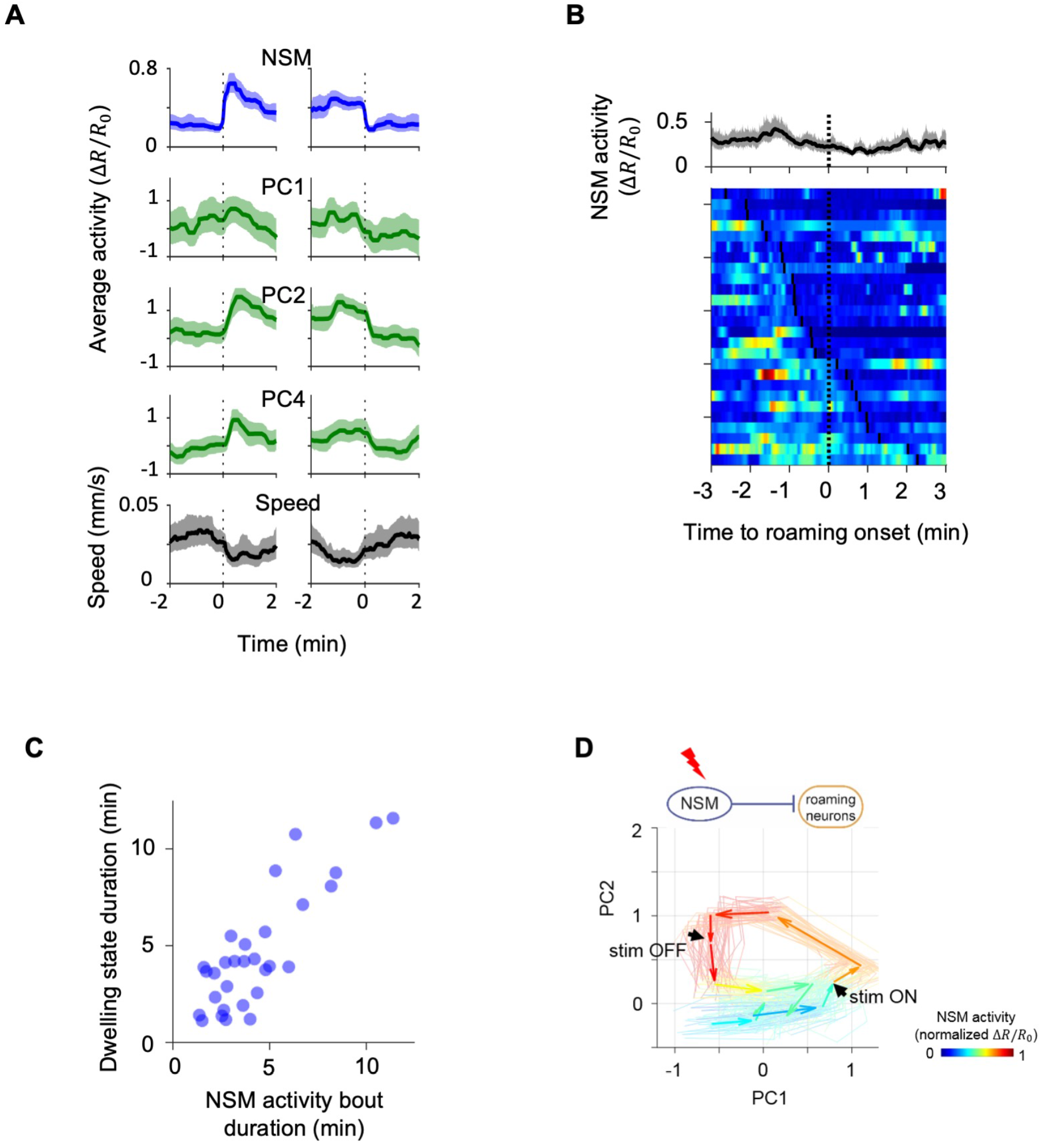
NSM activation coincides with dwelling states and optogenetic activation of NSM induces circuit activity changes akin to those seen during spontaneous state transitions. (A) Event-triggered averages centered on NSM activation (left) and termination (right) events. Data are from 17 animals and are shown as mean and 95% CI. PC4 is shown as an example to illustrate that dynamics beyond the first two principle components also change around the time of NSM activation. (B) NSM activity aligned to the onset of roaming states. (Top) Average NSM activity around the onset of roaming states. (Bottom) Heat map of NSM activity around instances of dwelling-to-roaming transition. Same color scale as in (D). Dotted black line denotes the onset of roaming states. Black ticks on the heat map mark the offset of an NSM activity bout. (C) Scatterplot of the durations of individual dwelling states and the durations of their coinciding NSM activity bouts. (D) Average trajectory of circuit activity in response to optogenetic activation of NSM, shown in principal component space. Each arrow represents average circuit dynamics over a 15 second interval. Color indicates ongoing NSM activity. Faint lines show bootstrap samples of the average dynamics. Black arrowheads indicate the beginning and the end of NSM::Chrimson activation.

**Figure 3 – Figure Supplement 1.**
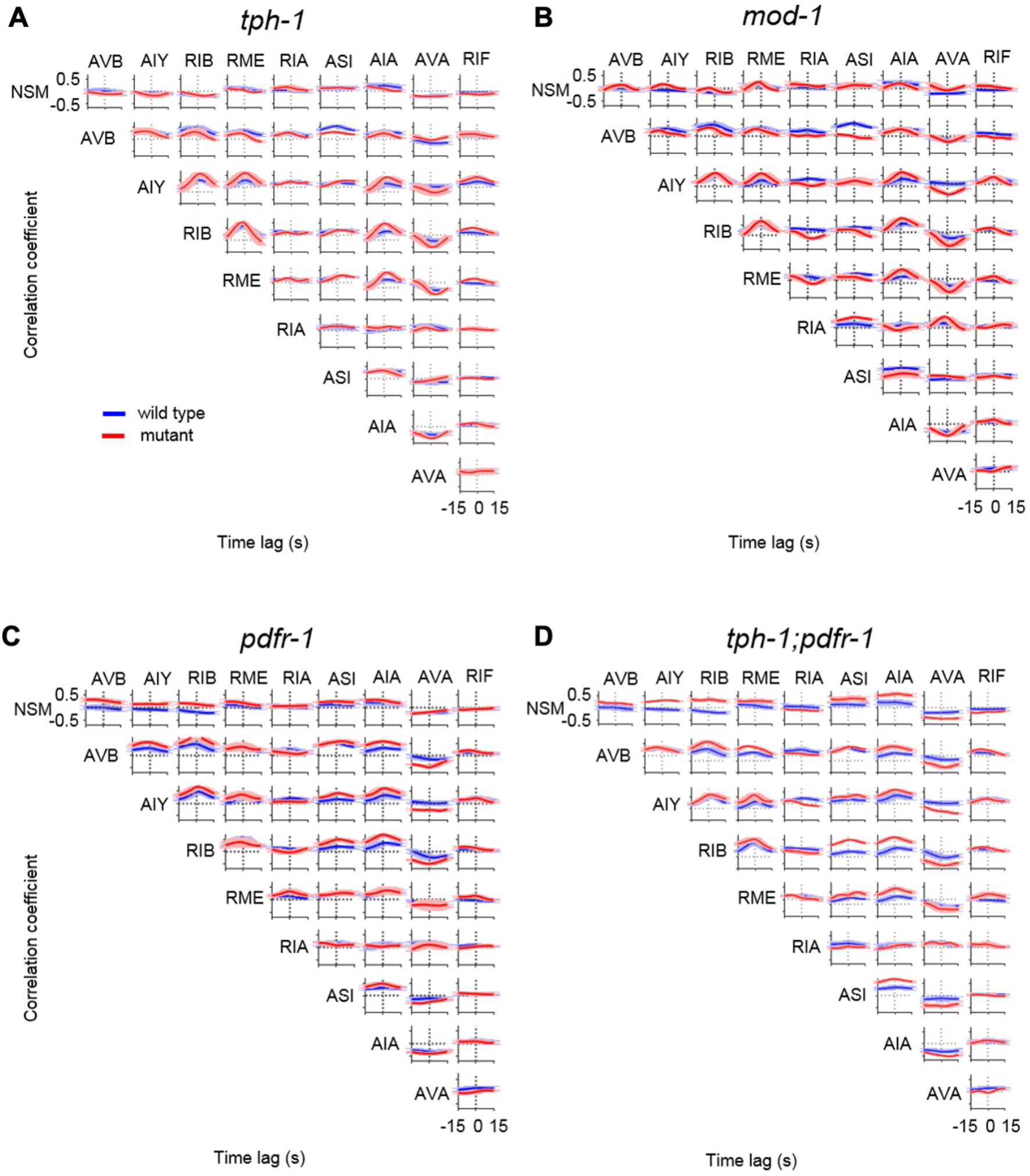
Cross-correlations between neurons in wild-type and mutant animals. (A-D) Average pair-wise cross-correlation function among neurons examined through freely-moving calcium imaging. Data are shown for the indicated genotypes as means and standard error. N = 17, 10, 8, 11, and 8 animals for WT, *tph-1*, and *mod-1*, *pdfr-1*, and *tph-1;pdfr-1* animals.

**Figure 3 – Figure Supplement 2.**
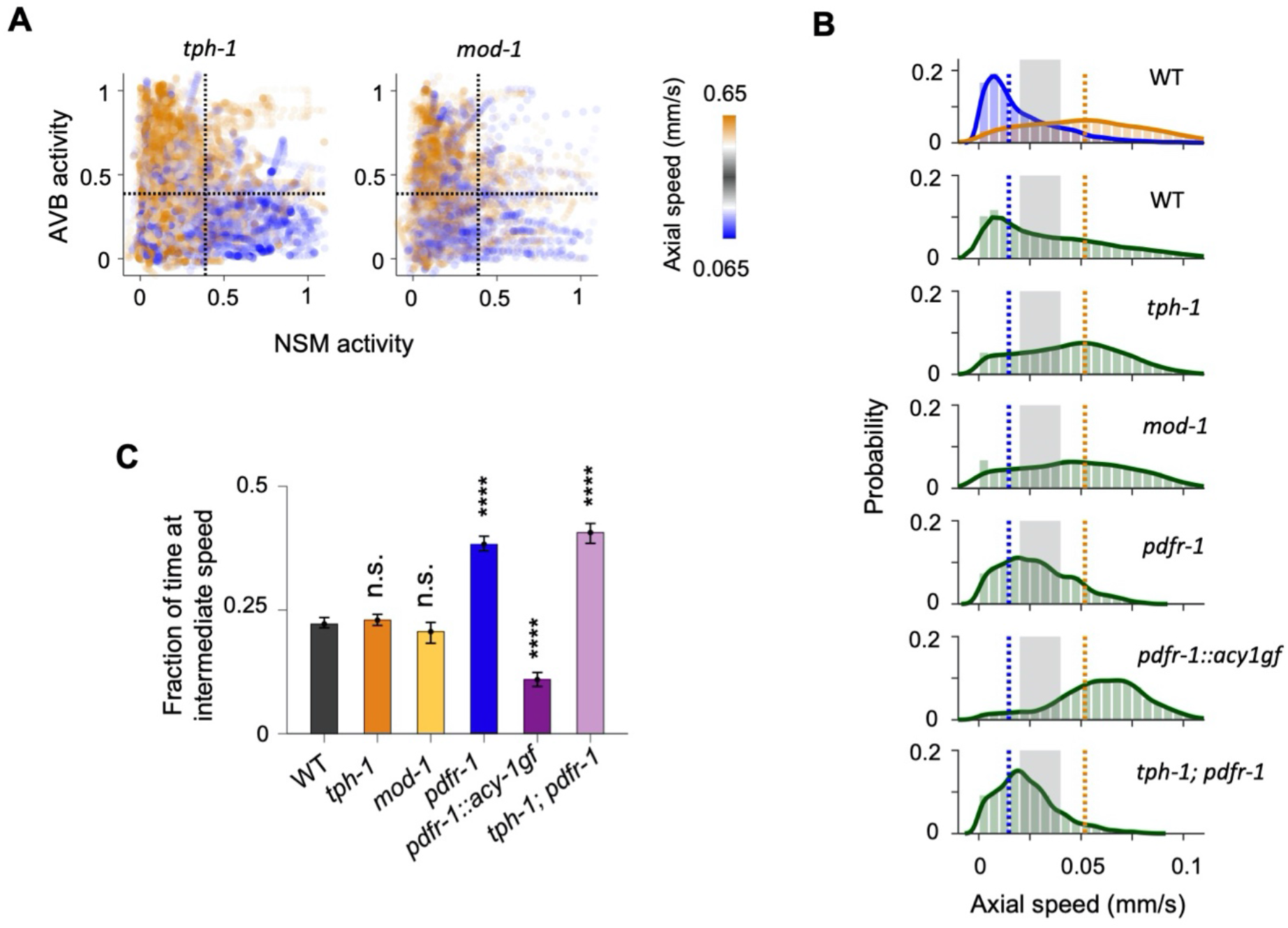
Analyses of circuit dynamics and foraging behavior in serotonin and PDF signaling mutants. (A) Scatterplots of NSM and AVB activity in *tph-1* mutants and *mod-1* double mutants. Data points are colored by the instantaneous speed of the animal. (B) Probability distribution functions of the axial speed of the animal in wild type and various 5-HT and PDF signaling mutants. Top panel shows speed distributions specific to the dwelling (blue) or the roaming (orange) states. Dotted blue and orange lines indicate the median speeds for the dwelling and roaming states, respectively. Shaded region defines the intermediate speed range used for the analysis in panel C. (C) Fraction of time animals moved at speeds intermediate between typical dwelling and roaming speeds for wild-type and mutant animals. The range of intermediate speeds is defined by the shaded region shown in panel B. Data are shown as mean and standard error. N = 17, 10, 8, 11, 9 and 8 animals for WT, *tph-1*, and *mod-1*, *pdfr-1*, *pdfr-1::acy-1gf* and *tph-1;pdfr-1* animals. *p<0.05, **p<0.01, ****p<0.0001, comparison of bootstrap distributions with BH correction.

**Figure 4 – Figure Supplement 1.**
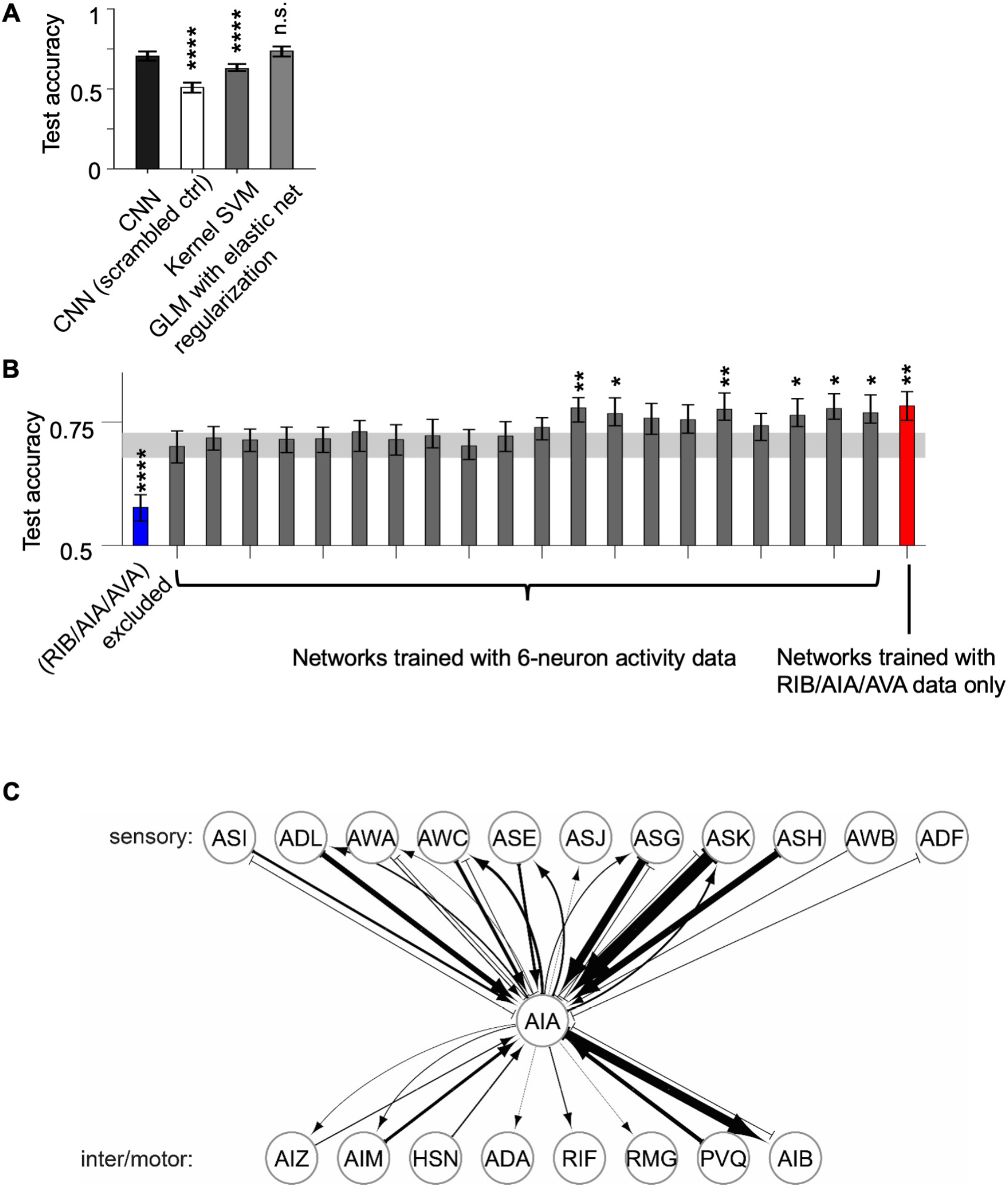
Evaluation of CNN classifier performance and connectivity diagram of AIA. (A) AUC-ROC of the CNN classifiers trained on authentic data compared to those trained on scrambled data and to the performance of two other common types of classifiers. Data are shown as mean and 95% CI from 200 training sessions. (B) AUC-ROC of CNN classifiers trained on data withholding different neuron triplets from the full data set, or with data from only the RIB, AIA, and AVA neurons. Gray band indicates the 95% CI of the accuracy of CNN classifiers trained on the full data set, as shown in A. (C) Synaptic inputs and outputs of the AIA neuron. Data are from the *C. elegans* connectome. Bilaterally symmetric pairs of neurons (e.g. AIAL and AIAR) were merged here for display purposes. Connections supported by only one single synapse were not included. Note the dense synaptic inputs onto AIA from chemosensory neurons.

**Figure 4 – Figure Supplement 2.**
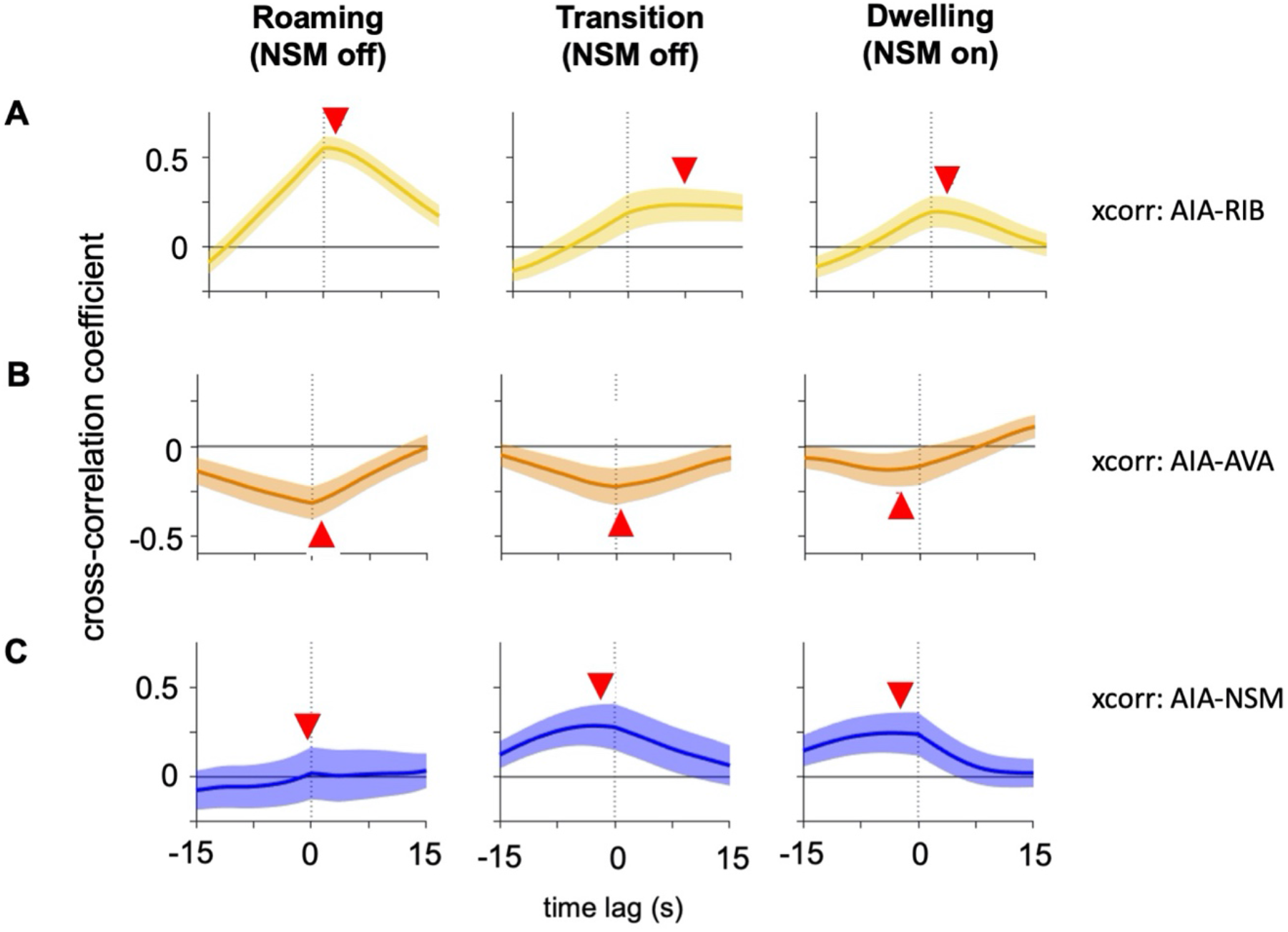
Activity correlations between AIA and other neurons during roaming and dwelling. (A) Average cross-correlation function between AIY and AIA activity at three time windows relative to NSM activation (from left to right): 1) 100 to 40.5 seconds prior, 2) 60 to 0.5 second prior, 3) 30 seconds prior to 30 seconds after (i.e. during NSM activity onset). (B) Average cross-correlation function between RIB and AIA activity during the same time windows as in (A). (C) Cross-correlation function between NSM and AIA activity during the same time windows as in (A). Red arrowheads denote the location of maximum absolute value of the average cross-correlation function. These values were used to generate the panels in Fig. 4F. Shaded regions denote 95% CI of the mean.

**Figure 6 – Figure Supplement 1.**
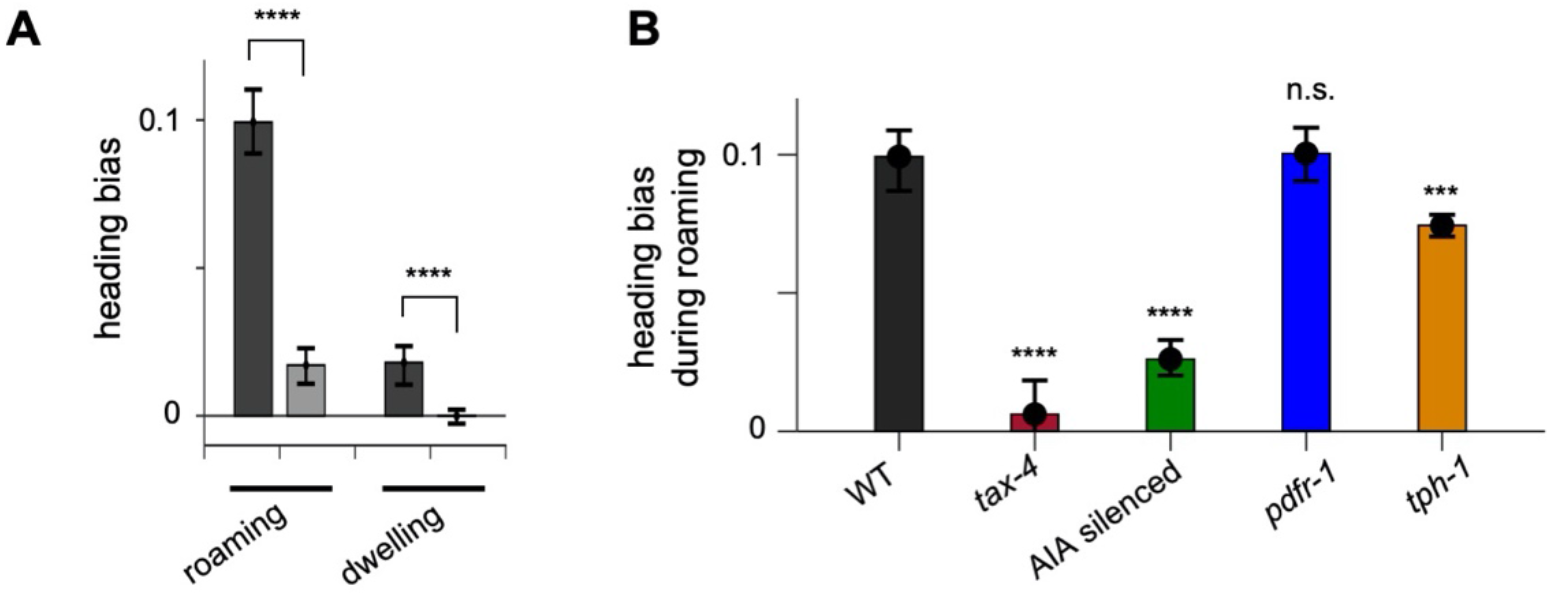
Food-directed navigation in patch foraging assays. (A) Heading bias during roaming versus dwelling in the patch foraging assay (dark bars) and control sparse food plates (light bars), shown separate for roaming and dwelling. (B) Heading bias during roaming for animals of the indicated genotypes. All error bars are 95% CI of the mean. ***p<0.001, ****p<0.0001, Wilcoxon rank sum test.

## Notes

### Competing Interest Statement

The authors have declared no competing interest.

